# Insects evolved a monomeric histone-fold domain in the CENP-T protein family

**DOI:** 10.1101/2024.11.15.623767

**Authors:** Sundar Ram Sankaranarayanan, Jonathan Ulmer, Anna Mørch, Ahmad Ali-Ahmad, Nikolina Sekulić, Ines Anna Drinnenberg

## Abstract

The histone fold domain (HFD) is a conserved protein interaction module that requires stabilization through a handshake interaction with an HFD partner. All HFD proteins known to date form obligate dimers to shield the extensive hydrophobic residues along the HFD. Here, we find that the lepidopteran kinetochore protein CENP-T is soluble as a monomer. We attribute this stability to a structural rearrangement, which leads to the repositioning of the HFD helix α3. This brings a conserved two-helical extension closer to the histone fold, where it takes over the position and function of the CENP-T partner CENP-W. This change has no effect on the DNA binding ability of the lepidopteran CENP-T. Our analysis suggests that the monomeric HFD originated in the last common ancestor of insects, with a possible second independent origin in acariformes, both of which lack CENP-W. Our study highlights an unexpected structural variation in a protein module as conserved and optimized as the HFD providing a unique perspective on the evolution of protein structure and the forces driving it.

## Introduction

The histone fold domain (HFD), a defining feature of histone proteins, is one of the most abundant and conserved DNA binding domains detected across eukaryotes and archaea and it is present in certain prokaryotic proteins (Talbert *et al*, 2019; Makarova *et al*, 2005; Postberg *et al*, 2010; Hocher *et al*, 2023; Henneman *et al*, 2018). The HFD is characterized by the presence of a long central helix flanked by two shorter helices that are connected by loops (Arents *et al*, 1991). The positively charged sidechains on the surface of the folded HFD favour interactions with the DNA backbone (Arents & Moudrianakis, 1995). The presence of exposed hydrophobic residues along the helices of the HFD renders HFD proteins unstable as monomers (Karantza *et al*, 1995, 1996; Banks & Gloss, 2004). A handshake-like interaction with a second HFD partner is required to shield these residues and stabilize the protein. Thus, the HFD can also be considered a protein-dimerization motif in addition to its role in DNA binding (Arents *et al*, 1991; Arents & Moudrianakis, 1995). Besides histones, the HFD is also commonly detected in proteins involved in transcription, DNA replication and repair, chromatin remodelling, as well as at the kinetochore (Figure 1A).

**Figure 1.**
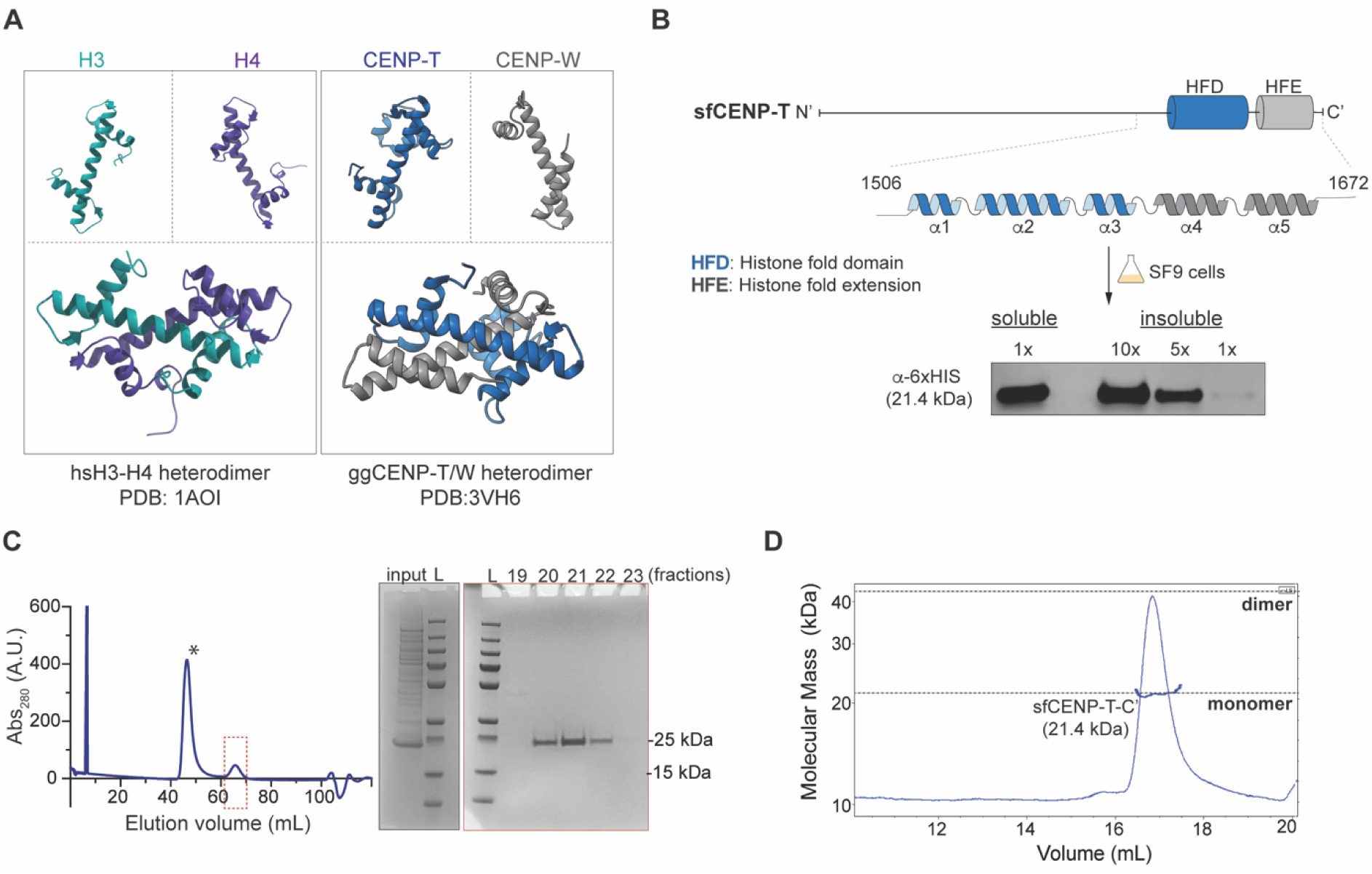
Lepidopteran CENP-T is a soluble monomeric protein. (A) The organization of helices of a canonical HFD monomer and their physiological heterodimeric state illustrated from the known structures of histones H3-H4 from humans (PDB: 1AO1) and CENP-TW from chicken (PDB:3vh6). (B) (*Top*) Line diagram of sfCENP-T with the relative positions of the Histone fold domain (HFD) and the histone fold extension (HFE) marked by blue and grey cylinders, respectively. The linear structure of each of these domains is expanded. (*Bottom*) Analysis of soluble and insoluble fractions of SF9 cells expressing the C-terminal fragment of 6xHis-sfCENP-T^1147-1314^ by western blot analysis. (C) The elute from affinity purification step for 6xHis-sfCENP-T^1147-1314^ fragment was further purified to homogeneity by size exclusion chromatography (SEC). The plot depicts the SEC elution profile with the *x-* and *y-* axes indicating elution volume and absorbance at 280 nm, respectively. The peak fractions marked by an asterisk and a dotted red box were visualized by SDS-PAGE stained with Coomassie Blue (also see Figure S1A). Input: a fraction of the elute from affinity purification step for 6xHis-sfCENP- T^1147-1314^ fragment loaded into the SEC column. L: molecular weight marker. (D) Multi-angle Light Scattering coupled with SEC (SEC-MALS) of the 6xHis-sfCENP-T^1147-1314^ fragment reveals the monomeric state of the protein (21.2 kDa). The molecular weight inferred from MALS is represented as a blue line overlaid on the SEC peak for this protein.

The kinetochore is a multiprotein complex assembled on the centromere of each chromosome that connects chromosomes to spindle microtubules, enabling accurate chromosome segregation during cell division. The kinetochore has two primary layers: an inner kinetochore called Constitutively Centromere Associated Network (CCAN) which is proximal to centromeric DNA, and an outer kinetochore that attaches to the spindle microtubules (Cheeseman, 2014; Ariyoshi & Fukagawa, 2023)(Cheeseman, 2014; Ariyoshi & Fukagawa, 2023). Two components of the CCAN, CENP-C and CENP-T, are critical to connect the inner to the outer kinetochore. CENP-C directly binds to nucleosomes containing a specialized histone H3 variant CENP-A (Falk *et al*, 2015; Kato *et al*, 2013; Carroll *et al*, 2010)that epigenetically marks centromeric chromatin and initiates kinetochore assembly in various eukaryotes. Via its N-terminus CENP-C, in turn, interacts with components of the outer kinetochore network (Milks *et al*, 2009; Przewloka *et al*, 2011; Screpanti *et al*, 2011). CENP-T binds DNA directly via a C-terminal HFD in the context of a CENP-TWSX nucleosome-like complex (Nishino *et al*, 2012; Hori et al, 2008). This complex partially wraps and super-coils linker DNA, thereby sharing structural and functional properties of histones (Yatskevich *et al*, 2022; Takeuchi *et al*, 2014). In addition, CENP-T also contains N-terminal motifs that bind the outer kinetochore proteins Ndc80 and Mis12 (Hori *et al*, 2008; Nishino *et al*, 2012; Malvezzi *et al*, 2013; Huis In ’t Veld *et al*, 2016). Tethering CENP-T to ectopic sites on the chromosome was shown to be sufficient to recruit outer kinetochore proteins and segregate minichromosomes (Hori *et al*, 2013; Schleiffer *et al*, 2012). The centromere localization and function of CENP-T is dependent on its heterodimerization with CENP-W, typical of HFD proteins (Schleiffer *et al*, 2012; Hori *et al*, 2008). The strong interdependence of CENP-T and CENP-W is also reflected in their simultaneous presence or absence across the tree of life (Tromer *et al*, 2019). The CENP-TW heterodimer oligomerizes with the CENP-SX heterodimer to form the nucleosome like structure that preferably binds to a ∼100 bp linker DNA in the presence of nucleosomes (Nishino *et al*, 2012; Takeuchi *et al*, 2014). It should be noted that tetramerization is not essential for CENP-T function at the centromere as CENP-T localization was not perturbed by CENP-S depletion (Amano *et al*, 2009; Nishino *et al*, 2012). However, mutations affecting the interactions between CENP-TW and DNA resulted in the loss of its kinetochore localization and defective mitosis (Nishino *et al*, 2012). This makes the HFD and its interactions with DNA a critical node in vertebrate kinetochore assembly. Besides DNA binding, CENP-T also interacts with the CENP-HIKM complex, additional CCAN components that contribute to the inner kinetochore assembly (Basilico *et al*, 2014; McKinley *et al*, 2015; Pekgöz Altunkaya *et al*, 2016). The interface between these two complexes was resolved to be a three-helix bundle formed by the two conserved helices of the CENP-T- Histone Fold Extension (HFE or two-helical extension) and an N-terminal helix from CENP-K (Pekgöz Altunkaya *et al*, 2016; Hinshaw & Harrison, 2020; Zhang *et al*, 2020). The interaction between CENP-T and the CENP-HIKM complex is crucial for kinetochore stability and function (Pekgoz Altunkaya *et al*, 2016; Basilico *et al*, 2014; McKinley *et al*, 2015).

In previous studies, we conducted proteomic analyses (IP/MS) combined with remote homology predictions to get insights into the composition and assembly of kinetochores in Lepidoptera (Cortes-Silva *et al*, 2020). Lepidoptera are interesting because they have lost the otherwise conserved centromere marker protein CENP-A and outer kinetochore linker protein CENP-C (Drinnenberg *et al*, 2014). These analyses uncovered the presence of orthologs of several kinetochore components common in other eukaryotes, including a divergent homolog of CENP-T. Using *Bombyx mori* as an experimental model system, we found that much like the human CENP-T homolog, *B. mori* CENP-T (bmCENP-T) is essential for viability. Depleting bmCENP-T disrupted the localization of other kinetochore subunits, resulting in mitotic defects. Tethering bmCENP-T to ectopic sites recruited outer kinetochore proteins like the Ndc80 and Mis12, as observed in yeast and chicken. Despite the functional conservation, we were unable to detect a potential homolog of the stabilizing partner CENP-W in Lepidoptera by homology searches or by affinity purifications of CENP-T or other conserved CCAN subunits (Cortes-Silva et al., 2020).

After observing the lack of CENP-W we aimed to address how its loss has been compensated. By analysing the structure of bmCENP-T, we report unprecedented changes in the HFD that alleviates the need for stabilization by an interacting partner. We detect a reorientation of α3 in the HFD that brings the helices of the HFE closer to the HFD, so that they occupy the position of CENP-W. In this arrangement, the extension might stabilize the hydrophobic residues of the HFD, thereby acquiring the function of CENP-W, while still retaining its conserved role in interacting with CENP-HIKM. Interestingly, this structural rearrangement of the HFD was present in CENP-T from all insect orders that lost CENP-W. In line with these observations, we find lepidopteran CENP-T to be a stable monomer in solution that retained its ability to bind DNA without the need of an interacting partner.

## Results

### Lepidopteran CENP-T is a soluble monomeric protein independent of CENP-W

To discover potential HFD partners of CENP-T that we might have missed in our previous analyses, we ectopically expressed the C’ terminal fragment of *Spodoptera frugiperda* CENP-T (sfCENP-T) that includes the HFD and the two-helical histone fold extension (HFE) in SF9 cells, a lepidopteran cell line derived from *Spodoptera frugiperda*. We detected the sfCENP-T fragment mainly in the supernatant (soluble) fraction of the cell lysate after centrifugation by western blot analysis (Figure 1B). The insoluble pellet fraction had relatively lower levels of the protein. For HFD proteins to remain soluble, they must form dimers—either by homo-dimerizing with themselves, as seen in archaeal histones (Stevens et al, 2020; Hocher & Warnecke, 2024; Mattiroli et al, 2017)(Stevens et al, 2020; Hocher & Warnecke, 2024; Mattiroli et al, 2017) or by heterodimerizing with another HFD protein, as it is the case for the vertebrate CENP-T and CENP-W. To distinguish between these two possibilities, we purified the sfCENP-T fragment over size-exclusion chromatography (SEC) and performed Multi Angle Light Scattering (MALS) analyses to determine the molecular weight of the potential complex that it might form. Analysis of the fractions enriched for the purified protein by SDS-PAGE did not reveal any coeluting proteins (Figures 1C and S1A) supporting the absence of a previously undetectable ortholog of CENP-W or the formation of a complex with another histone fold proteins. SEC-MALS analyses further revealed that the sfCENP-T fragment was a monomer in solution, hitherto unknown for any HFD-containing protein (Figure 1D). We also detected the same sfCENP-T fragment in the soluble fraction when expressed in a heterologous host like *E. coli* (Figure S1B), thereby corroborating that the HFD in lepidopteran CENP-T folds independent of an interacting partner and is a monomer in solution.

### An acquired role of the histone fold extension in stabilizing CENP-T

To understand the molecular basis of this stability, we used AlphaFold to model the structure of full-length CENP-T from *Bombyx mori*, the most established lepidopteran model organism in which CENP-T was functionally characterized (Figure 2A). The C-terminus with the HFD and HFE were highly similar across different AlphaFold models and were generated with high confidence (pLDDT>90) (Figure S2A-B). We also modelled the sfCENP-T which showed a high structural similarity to the predicted bmCENP-T structure (Figure S2C). A comparison of the best-ranked model of bmCENP-T with the structure of the canonical CENP-TW heterodimer (as reported for chicken) revealed two striking rearrangements. First, the third helix (α3) of the HFD is positioned parallel to the second helix (α2) in bmCENP-T (that we refer to as cryptic HFD hereafter) as opposed to its position perpendicular to the α2 in the conventional HFD. Second, this altered position of α3 enables the HFE to be closer to the HFD compared to the canonical CENP-T (Figure 2A-B).

**Figure 2.**
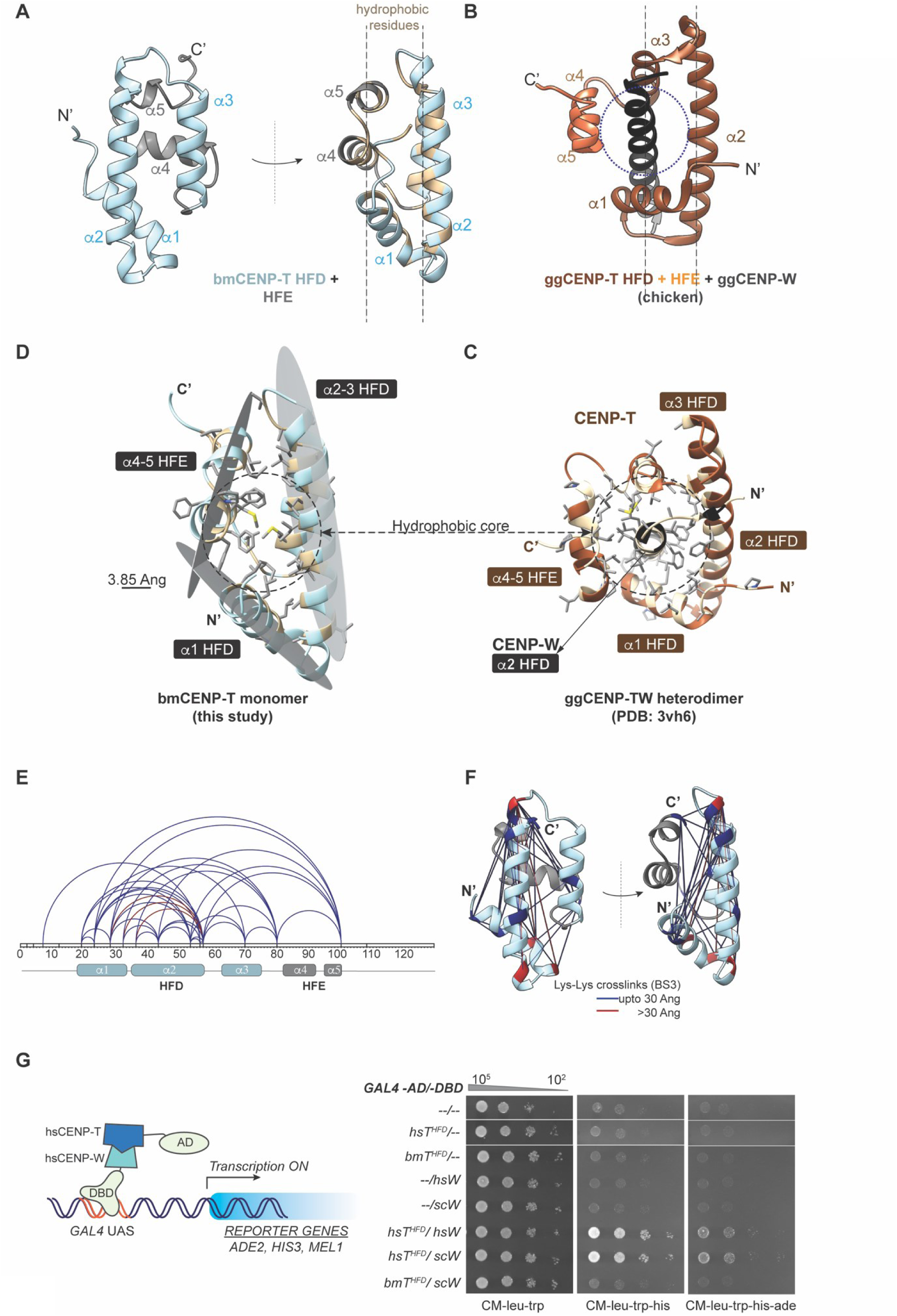
The cryptic histone fold and the repositioned extension occludes CENP-W and stabilizes CENP-T as a monomer. (A) Front and side views of the AlphaFold model generated for the bmCENP-T fragment containing the HFD and HFE. Hydrophobic residues within this fragment are coloured beige in the side view. (B) A comparative side view of ggCENP-TW heterodimer highlighting the position of ggCENP-T-HFE relative to ggCENP-W and the ggCENP-T-HFD (dotted circle). For clarity, only the second helix of ggCENP-W histone fold is shown in black. (C) Illustration of the proposed stabilization of bmCENP-T histone fold by hydrophobic residues present in the HFE. Three hydrophobic surfaces from α1, α2-3, and α4-5 are proposed to form a hydrophobic core. The hydrophobic residues and their side chains are coloured as in (A). Scale bar, 3.85 Ang. (D) A tilted view of ggCENP-TW complex shown in panel B depicting the proximity of the sidechains of hydrophobic residues from the helices of ggCENP-T and ggCENP-W. The hydrophobic residues are shaded beige in the backbone and their sidechains are represented as sticks in grey. (E, F) BS3 crosslinking mass spectrometry of bmCENP-T fragment represented in a 1D plot (E) and in the 3D model (F). Lysine pairs that are crosslinked in the experiment and are within 30 Å in the AlphaFold model (distance cutoff for BS3) are connected by blue lines. The crosslinked lysine pairs that exceed this distance cutoff in the AlphaFold model are shaded and connected in red. See also Table S1. (G) Schematic representation of a yeast two-hybrid assay (*left panel*). The transcription of reporter genes is driven by the interaction of proteins that are fused to the GAL4-AD and GAL4-DBD proteins respectively. The known interacting proteins hsCENP-T and hsCENP-W are shown as representative examples in this schematic. Results from spot dilution assay performed with the reporter strain transformed with plasmids encoding the indicated GAL4-AD and -DBD fusions is shown in the *right panel*. Growth on CM-leu-trp serves as control. Growth on CM-leu-trp-his and CM-leu-trp-his-ade dropout plates indicate positive interaction between the proteins cloned in fusion with GAL4-AD and GAL4-DBD. The plates were photographed after incubation at 30°C for 48 hours.

The position of α3 in the canonical fold allows access to CENP-W such that it is trapped between the helices of the HFD and the extension. This configuration creates two sets of hydrophobic surfaces - the central helix of CENP-W and the buried part of the helices of HFD and the extension. This arrangement facilitates the formation of a hydrophobic core that stabilizes the heterodimer (Figure 2C). In the case of bmCENP-T, the rearrangements in the cryptic HFD brings the two helices of the HFE in the position otherwise occupied by CENP-W. The proximity between the hydrophobic residues of the HFE and those of α1-2 of the HFD enables the formation of a hydrophobic core, making the protein fragment stable and soluble independently of CENP-W (Figure 2D).

To confirm the AlphaFold predictions, we initially attempted to solve the structure of the BmCENP-T HFD by crystallography, but we were unsuccessful (see *supplemental information*). We therefore used Crosslinking Mass Spectrometry (CLMS) to identify intra-protein contacts and assessed the distance between these pairs in the AlphaFold model. More than 90% of the crosslinks detected in our experiment were within the 30 Å reach of the BS3 crosslinker, indicating an overall fit of the AlphaFold model to the experimental data (Figure 2E-F, Table S1). This further increases our confidence in the predicted structure. Overall, these results provide an explanation for the biochemical behaviour of the monomeric bmCENP-T fragment, which involves a role for the two-helical CENP-T extension in shielding the hydrophobic interface of the histone fold domain.

### The cryptic HFD and repositioned extension occludes the interaction with CENP-W

From the structural predictions of bmCENP-T, we inferred that the HFE occupies the channel conventionally occupied by CENP-W. To further explore the impact of the altered organisation of the bmCENP-T C-terminus on its interaction with CENP-W, we used a yeast two hybrid assay (Figure 2G). We used the interaction between the human CENP-T C-terminus (HFD and the extension) and human CENP-W as positive control, because both proteins have a canonical HFD and are known interacting partners (Figure 2G). We were also able to detect the interaction between the canonical HFD pairs human CENP-T and *S. cerevisiae* CENP-W (scCENP-W), suggesting that the assay was sensitive enough to detect interaction between proteins with compatible folds but from phylogenetically distant hosts (Figure 2G). However, we could not detect an interaction between the bmCENP-T C-terminus (cryptic HFD and extension) with scCENP-W (Figure 2G, S2D). AlphaFold Multimer predictions also suggested an incompatibility between bmCENP-T and scCENP-W (Figure S2E). Collectively, these observations concur with our model that the structural rearrangements in bmCENP-T prevent its interaction with CENP-W.

### The DNA binding ability of CENP-T is conserved in *B. mori*

Given the divergence of *B. mori* CENP-T from the canonical CENP-T structure, we next tested if the DNA binding ability of *B. mori* CENP-T was retained using electrophoretic mobility shift assays. Like the canonical CENP-TW complex (Nishino *et al*, 2012), the bmCENP-T C-terminal fragment [bmCENP-T (wt)] was able to shift DNA, suggesting that the cryptic HFD retained the DNA binding ability. We observe maximum DNA-protein complex formation by incubating a 12 bp template DNA with 5x-molar excess of protein (Figure 3A, S3A).

**Figure 3.**
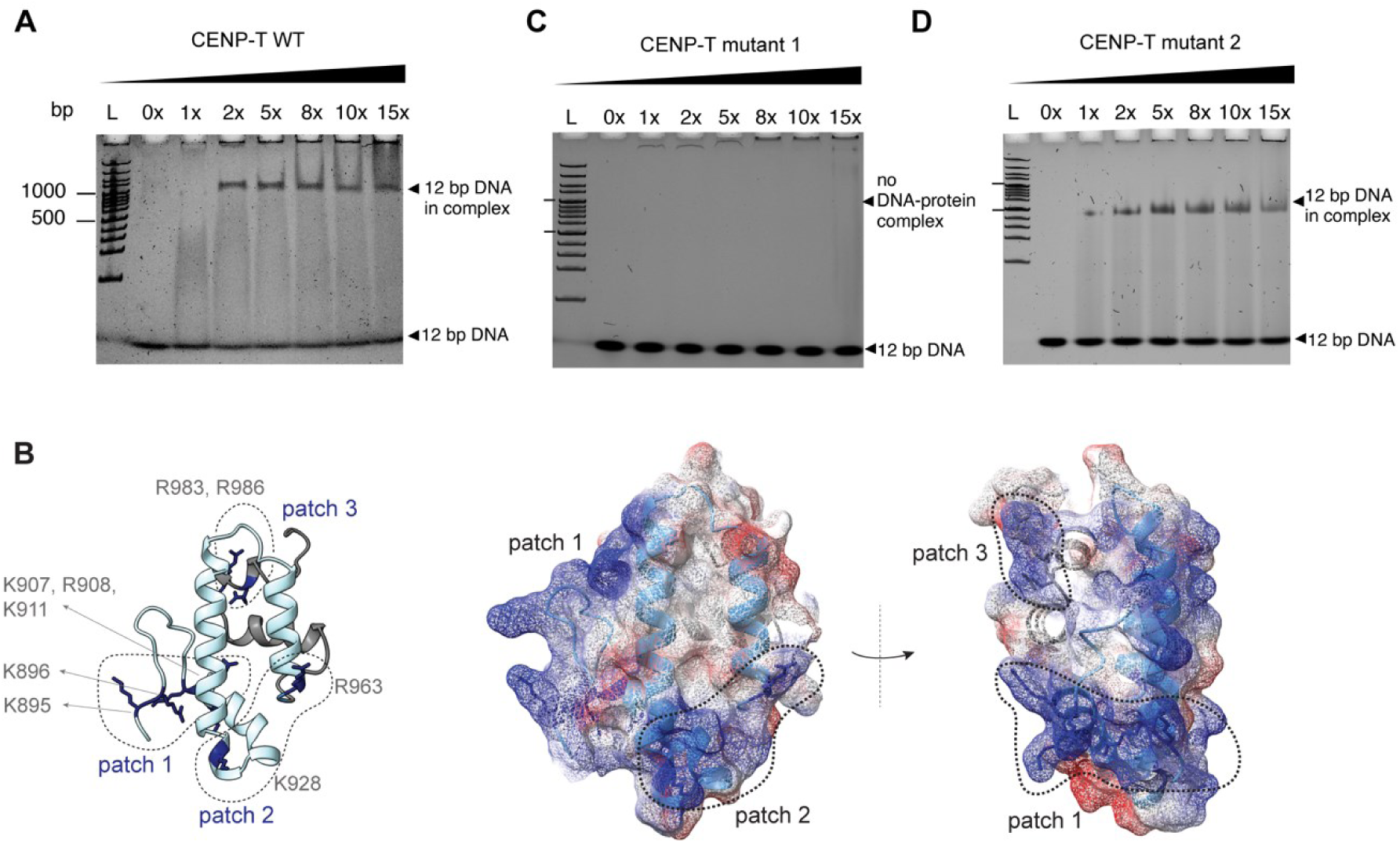
The DNA binding ability of CENP-T is conserved in *B. mori*. (A) 5% Native PAGE gels of DNA titrations using different molar ratios of bmCENP-T^894-1016^-WT with a 12 bp DNA in 100mM NaCl. Lane L: 100 bp DNA marker. (B) The positively charged surfaces formed by the conserved lysines and arginine residues in patches 1-3 are highlighted. The position of these residues and their sidechains (dark blue) is shown in the ribbon model in the left. The surface charge distribution of the same model is depicted in right. The front and side views are shown for this model with the positively and negatively charges surface coloured blue and red respectively. (C-D) Native PAGE gels of DNA titration using different molar ratios of bmCENP-T^894-1016^ patch 1 mutant with the substitutions K895A, R896A, K907A, R908S, and K911A (in C) and bmCENP-T^894-1016^ patch 2 mutant with the substitutions R928S and R963S (in D). The assay was performed with a 12 bp DNA fragment and indicated amounts of protein at 100 mM NaCl. Lane L: 100 bp DNA marker.

To further dissect the regions of bmCENP-T that bind to DNA, we identified positively charged regions on the protein surface (Figure 3B). By comparing these with the residues known to contact DNA in human CENP-T (Nishino et al., 2012), we assigned three patches that might mediate DNA interactions along the bmCENP-T fragment, namely patch 1 (located at α1 and sequences upstream of it, with residues K895, R896, K907, R908, K911), patch 2 (with residues K928, R963 located across α2-3), and patch 3 (with residues R983, R986 located at the C’ extension) (Figure 3B) These residues show a high degree of conservation across lepidopteran CENP-T sequences (Figure S3B). To assess the relative contribution of each of these patches in DNA binding, we mutated all Arg and Lys residues in each of these patches to Ala and Ser, respectively. The SEC elution profile of bmCENP- T^patch1^ and bmCENP-T^patch2^ mutant proteins were similar to the bmCENP-T indicating that these mutations do not affect the overall folding of the protein (Figure S3C). Early elution of CENP-T^patch3^ mutant in the SEC indicated deviation from the WT structure and hence this protein was not considered for subsequent analysis.

Among the mutants tested, we observed a loss in DNA binding only for the bmCENP-T^patch1^ mutant suggesting a critical role for patch 1 residues in contacting DNA (Figure 3C). bmCENP-T^patch2^ mutations had no effect on DNA binding (Figure 3D). To dissect the DNA-patch 1 interactions further, we sub-categorized these residues into patch 1.1, which includes residues upstream of α1 (K895, R896) and patch 1.2, which includes residues within α1 (K907, R908, K911). Both mutants failed to bind DNA showing that the entire patch 1 plays an essential role in binding DNA (Figure S3E-F). These studies demonstrate that the DNA binding function of CENP-T is conserved despite the changes in the histone fold.

### The cryptic HFD can be traced to the last common ancestor of insects

Having demonstrated that lepidopteran CENP-T can fold stably and associate with DNA in the absence of CENP- W, we expanded the analysis of CENP-T structures to other insect orders. With the exception of species belonging to the order Coleoptera (beetles), CENP-T homologs are present in all other insect orders. However, no insect CENP-W homolog has been identified to date. Based on the AlphaFold predictions of CENP-T structures of each insect order, the cryptic histone fold is present in all insect orders including Ephemeroptera (mayflies), the earliest branching insect order (Figure 4). This conservation places the evolution of cryptic HFD to the last common ancestor of insects, which might have co-occurred with the loss of CENP-W.

**Figure 4.**
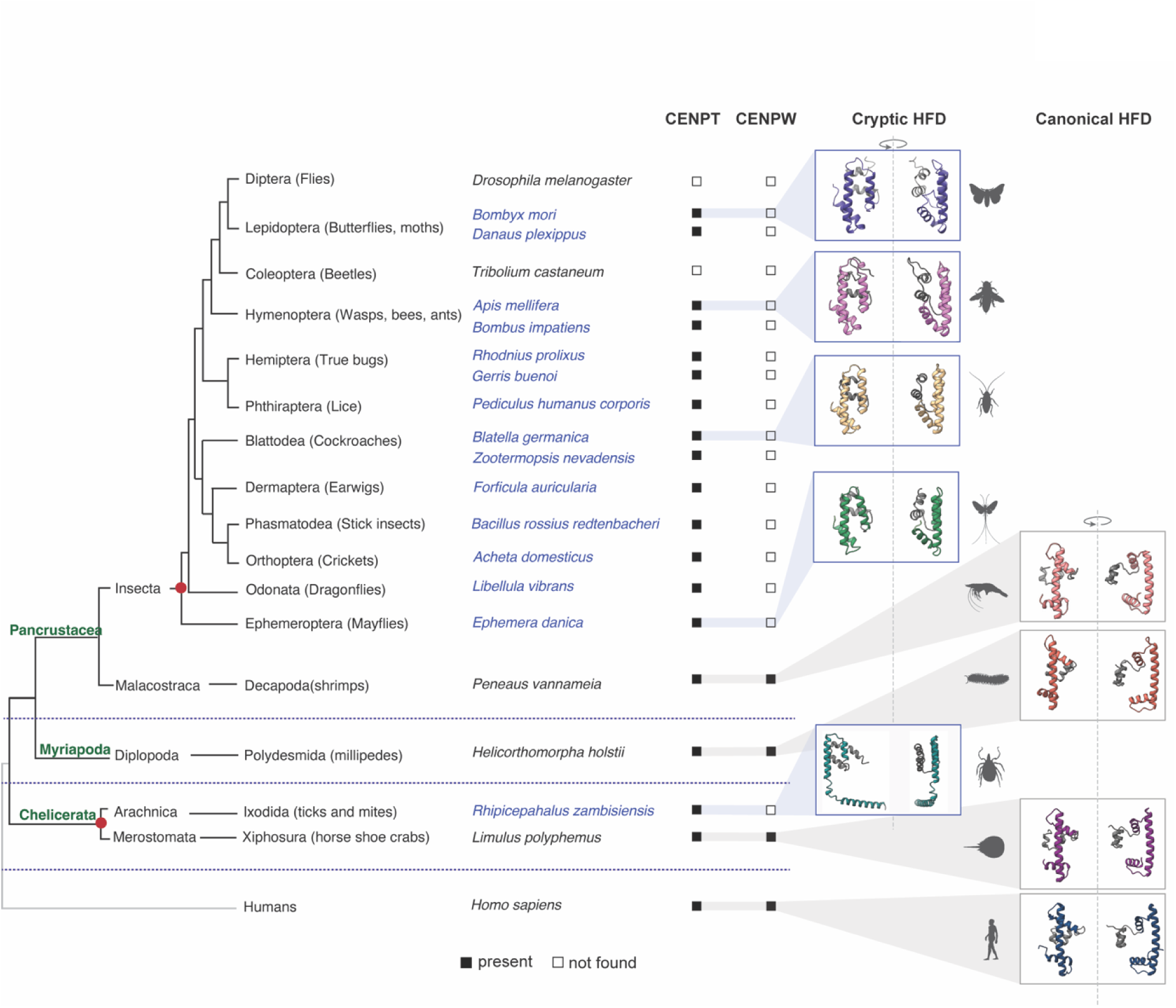
Independent evolution of the cryptic HFD in Arthropods. A cladogram depicting the three major arthropod classes Chelicerata, Myriapoda, and Pancrustacea, with humans as an outgroup. Species representative of each of these classes are mentioned along with their respective subclass and order. The presence/ absence of indicated proteins in each species is marked by filled and empty boxes, respectively. For protein IDs see Table S2. The configuration of the CENP-T HFD was assessed using AlphaFold for one species per each taxonomic order. Species with a cryptic CENP-T HFD are coloured in blue, and the ones with a canonical CENP-T HFD are coloured black. The cartoons on the right are two different views of the predicted model to highlight the proximity of the C’ extension. The red circles mark the two independent origins of the cryptic HFD identified in this study.

To understand the sequence variations associated with the evolution of this cryptic fold, we measured the amino acid substitutions across the HFD using Shannon’s diversity index. Due to the evolutionary conservation of HFDs, we expect a minimization of Shannon’s diversity score (entropy) for the residues in this domain, which we observed when conventional HFD sequences from vertebrates were analysed (Figure S4A). In the case of insects, we see reduction in sequence diversity only for the residues of α1 and the N’ half of α2. For residues corresponding to regions folded differently in the cryptic fold, we observe a marked increase in sequence diversity hinting at an ongoing evolutionary process or lower constraints on the amino acid identity in this region (Figure S4B).

We extended our search beyond insects to include other arthropod clades and were able to identify additional CENP-T homologs in species belonging to Xiphosura (ex: horseshoe crabs), Ixodida (ex: ticks), Polydesmida (ex: millipedes), and Decapoda (ex: shrimps). We generated AlphaFold models for each of these CENP-T homologs. In *Limulus polyphemus* (horseshoe crab), *Helicorthomorpha holstii* (millipede) and *Paneaus vannameia* (shrimp), CENP-T adopts a canonical HFD structure. Consistent with this, we also identified homologs of CENP-W in each of these organisms. In contrast, in the tick *Rhiphicephalus zambisiensis*, CENP-T was predicted to adopt the altered HFD structure with proximal HFE as described in this study (Figure 4). We failed to detect a CENP-W homolog in this species. Intriguingly, we could also detect other configurations of the HFD in related tick species where CENP-W homolog remains undetected. Notable example includes a configuration wherein the three helices of the HFD adopt a canonical structure, but the space occupied by CENP-W- α2 now hosts an alpha helix from the N’ of CENP-T itself (Figure S4C-E). Like the cryptic HFD, this configuration could also stabilize CENP-T to make it an independently folding protein. This might mark a second independent origin of a cryptic HFD within the arthropods, highlighting the adaptability of CENP-T proteins to bottlenecks like the loss of an interacting partner.

## Discussion

In this study, we report an unprecedented rearrangement in the structure of the HFD in insect CENP-T orthologs. This structural change alleviates the need of a HFD-partner and renders the protein soluble as a monomer. This is highly unusual for the family of HFD proteins known to be obligate dimers across the tree of life. While the HFD shows limited sequence conservation, both the tertiary structure and its functional organization as a dimer stabilized by a handshake interaction with an HFD-partner are conserved across the diverse range of proteins with this domain. The identification of a cryptic variant in such a highly conserved and optimized protein module offers a unique window to understand the evolution of protein structure.

We attribute the stability of monomeric CENP-T to the two-helical extension at the C-terminus, which is in proximity to the histone fold due to the repositioning of the α3 in the cryptic HFD as compared to a canonical CENP-T (Nishino *et al*, 2012). In canonical CENP-T, the HFD is critical for the interaction with CENP-W and DNA, while the two-helical extension is critical for the interaction with CENP-K of the CENP-HIKM complex (Hori *et al*, 2008; Nishino *et al*, 2012; Pesenti *et al*, 2022; Yatskevich *et al*, 2022; Yan *et al*, 2019; McKinley *et al*, 2015; Hinshaw & Harrison, 2020; Zhang *et al*, 2020). Despite the observed rearrangements in the HFD, bmCENP-T still binds DNA. This interaction is dependent on residues in the α1 of the HFD and the region upstream of it, both of which are also the regions of CENP-T that contact DNA in human and yeast CCAN/inner kinetochore complexes (Figure S3G) (Yan et al, 2019; Yatskevich et al, 2022; Pesenti et al, 2022). The sequence of the two-helical extension that mediates CENP-T recruitment to the kinetochores by interacting with the CENP-HIKM complex in vertebrates is also conserved in Lepidoptera. The loss of CENP-T localization upon depletion of CENP-I and CENP- M in *B. mori* cells suggests that the function of the two-helical extension is conserved as well (Cortes-Silva *et al*, 2020). We propose that the stabilization of the cryptic HFD is an acquired, second function of the two-helical extension in Lepidoptera. Taken together, our study highlights a structural innovation in CENP-T that support its stability even in the absence of CENP-W, while leaving the other functional associations of CENP-T unperturbed.

The presence of a C- terminal two-helical extension that is unique to CENP-T orthologs makes CENP-T uniquely poised among HFD proteins to evolve the ability to become soluble as a monomer. That said, the two-helical extension is highly conserved in CENP-T from yeast to humans irrespective of the structure of the HFD or the availability of CENP-W (Hinshaw & Harrison, 2020; Zhang *et al*, 2020; Cortes-Silva *et al*, 2020; Pekgöz Altunkaya *et al*, 2016). This leads us to hypothesize about the evolutionary drivers and constraints that may have triggered the origin of a cryptic fold in the last common ancestor of insects, and not in other lineages. The conserved HFD enables binding to CENP-W, which is necessary for the recruitment of CENP-SX as part of a tetrameric CENP-TWSX complex (Takeuchi *et al*, 2014; Nishino *et al*, 2012). While CENP-SX is not essential for cell viability, depletion of CENP-SX subunits in vertebrates leads to defective outer kinetochore formation and defects in mitotic progression (Amano *et al*, 2009; Nishino *et al*, 2012). The contributions of CENP-SX to outer kinetochore assembly might act as a functional constrain to retain these subunits at the kinetochore and in turn preserve a canonical CENP-T HFD in vertebrates and fungi. In insects, CENP-SX subunits are either lost or function mainly in DNA repair (Drinnenberg *et al*, 2014; Cortes-Silva *et al*, 2020). Their absence at the insect kinetochore renders the formation of a heterodimer between CENP-T and CENP-W redundant. Under these circumstances, the evolution of an altered histone fold might be advantageous because it removes the dependency on CENP-W. This optimization gains significance in Lepidopteran systems that lost the classical CENP-C based link between the inner and outer kinetochore and CENP-T remains to be the only direct linker (Cortes-Silva *et al*, 2020). Finally, the potential recurrence of the cryptic HFD of CENP-T in other arthropods raises the intriguing question whether similar changes might have occurred in additional eukaryotic lineages.

## Acknowledgements

SRS acknowledges the financial support from EMBO (LTF 505-2021). IAD receives salary support from the CNRS. Research in the Drinnenberg lab is supported by the Labex DEEP ANR-11-LABX-0044 part of the IDEX Idex PSL ANR-10-IDEX-0001-02 PSL, an ATIP-AVENIR Research grant, Institut Curie, the ERC (CENEVO-758757). Research in Sekulić lab is supported by the NCMM and the Research Council of Norway (grant numbers 187615 and 325528). We thank Gaétan Cornilleau from the Drinnenberg lab for generating the plasmids for the yeast two-hybrid experiment. We thank the EMBL proteomics platform for support with CLMS analysis, Ahmad El Marjou from the Curie recombinant protein platform, Carlos Kikuti from Institut Curie for help with the SEC-MALS analyses, and Vikram Alva-Kullanja (Max Plank Institute for Biology, Tubingen) for advice on evolution of the histone fold domain. We also acknowledge the support from Fabien Ferrage, Guillaume Bouvignies, and Theodore Bellon (Ecole Normale Supérieure - Département de Chimie, Paris) for help with preliminary NMR analysis. We thank Stanislau Yatskevitch (Genentech, USA) for critical reading of the manuscript. We would also like to thank Life Science Editors for editing services (www.lifescienceeditors.com).

---

**Figure S1.**
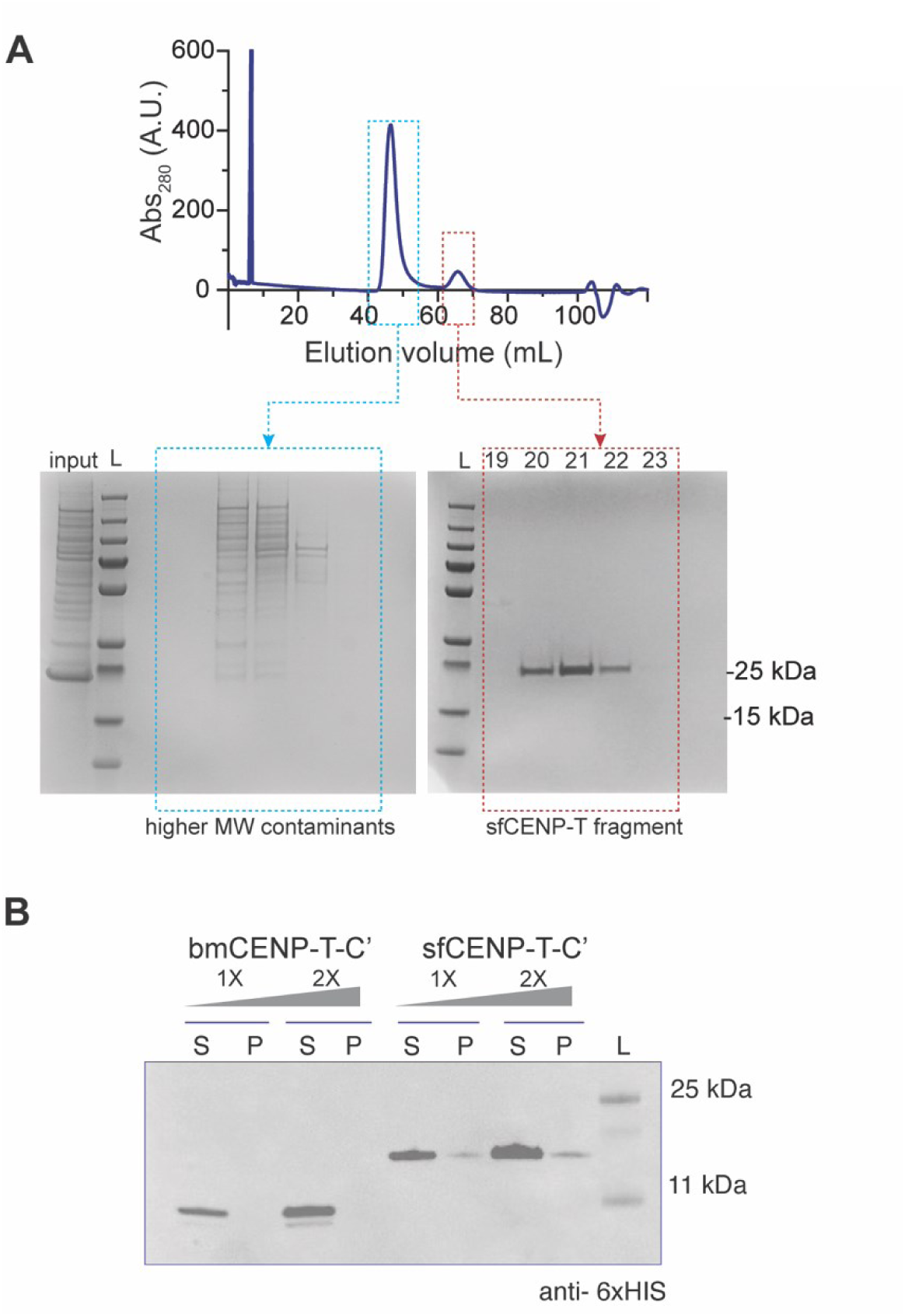
Solubility of lepidopteran histone fold in SF9 and *E. coli* cells. (A) SEC profile and the corresponding SDS-PAGE analysis of the indicated fractions of 6xHis-sfCENP-T^1147-1314^ purification. Input: a fraction of the sample loaded into the SEC column, L: molecular weight marker. (B) Western blot analysis of the soluble (S) and pellet (P) fractions of *E. coli* BL21 cells expressing His-bmCENP-T and His-sfCENP-T fragment with anti-His antibodies. L: molecular weight marker.

**Figure S2.**
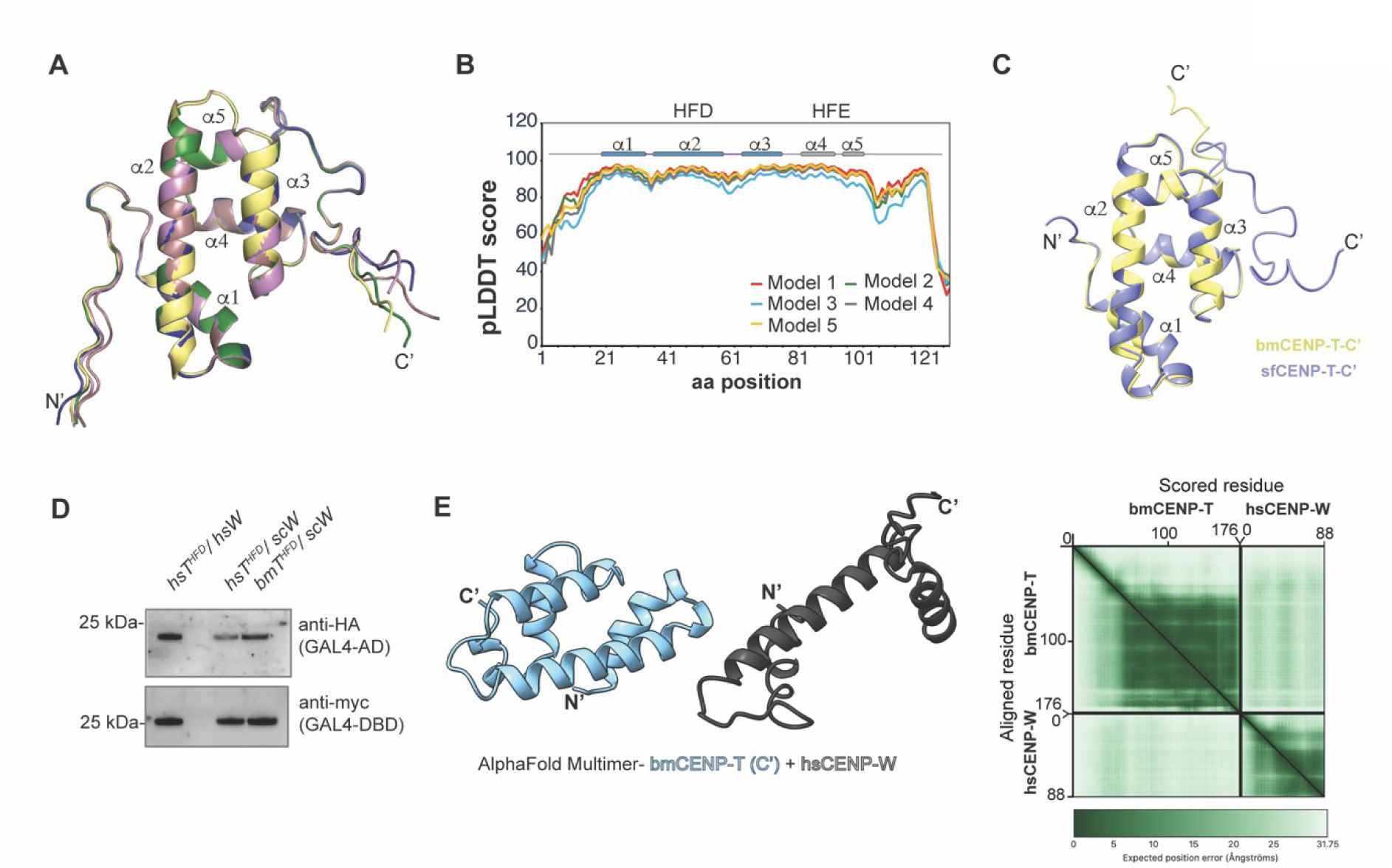
Structural basis for the stability of bmCENP-T in the absence of an interacting partner. (A) Overlay of the top five bmCENP-T models generated by AlphaFold. The helices of the HFD (α1-3) and the extension (α4-5) are indicated. (B) The pLDDT plot for the five models shown in (A). The regions corresponding to the helices of the HFD, and the extension are highlighted by a line diagram above the plot. (C) The best ranked AlphaFold models for bmCENP-T and sfCENP-T are overlaid to highlight structural similarities at the C-terminus. Only the regions corresponding to the HFD and the HFE are shown in the panel. (D) Whole cell extracts from cells used in the spot dilution assay shown in Figure 2G were analysed by immunoblotting using anti-Myc and anti-HA antibodies. (E) The inability of a cryptic HFD (from bmCENP-T) to interact with a canonical HFD (from hsCENP-W) as revealed by AlphaFold multimer. The predicted aligned error (PAE) matrix is presented on the right.

**Figure S3.**
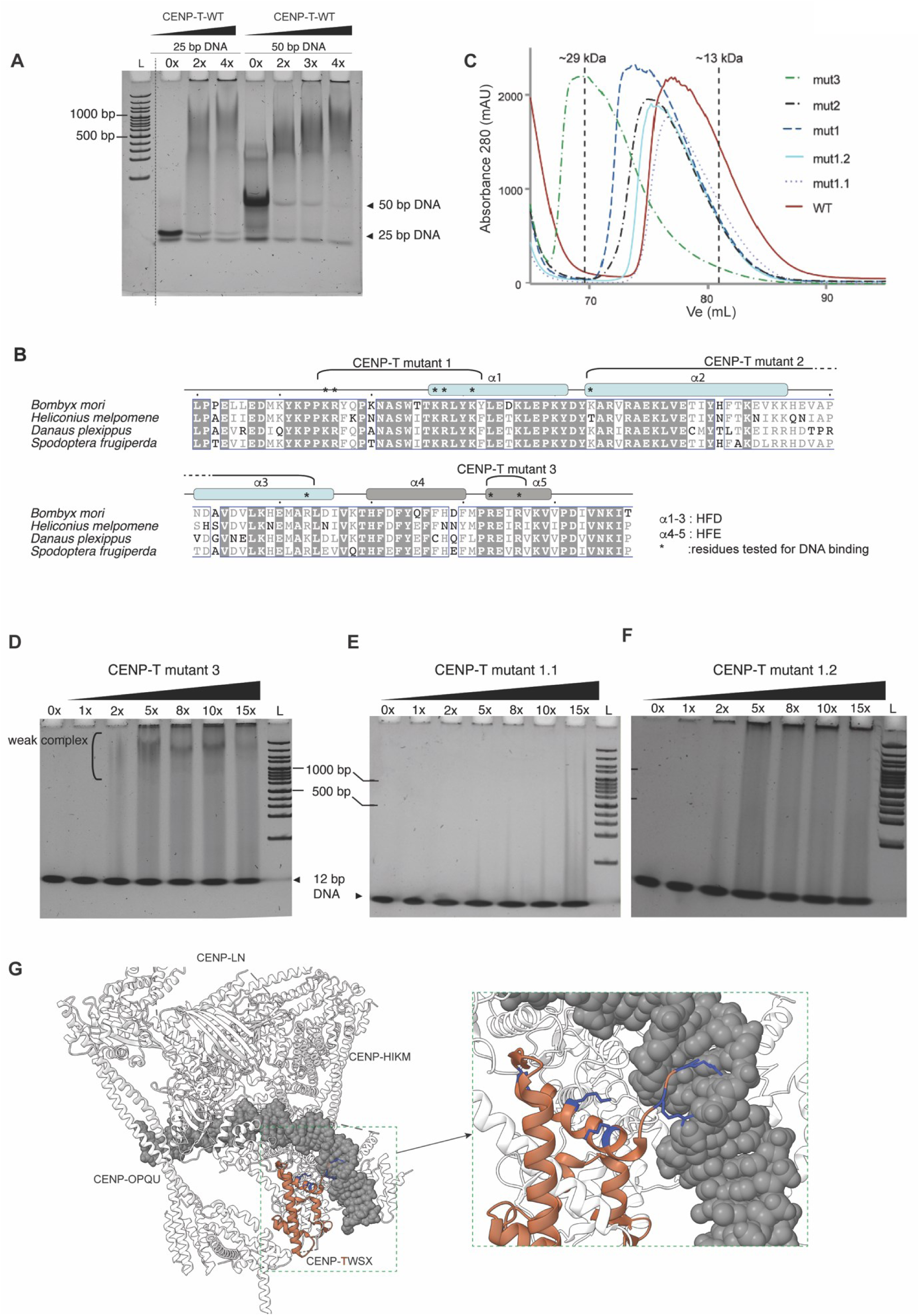
Residues in patch 2 and patch 3 are dispensable for DNA binding. (A) Native PAGE gels of DNA titration using different molar ratios of bmCENP-T^894-1016^-WT and a 25 bp or a 50 bp DNA fragment at 100 mM NaCl. Lane L: 100 bp DNA marker. (B) Multiple sequence alignment of the C’ fragment of CENP-T between 4 lepidopteran species. Identical residues are highlighted with a grey background. The residues tested for DNA binding function are marked with an asterisk. (C) The SEC elution profile of the WT and mutant versions of bmCENP-T used in the DNA binding assays in Figure 3 and Figure S3. (D-F) Native PAGE gels of DNA titration using different molar ratios of bm CENP-T^894-1016^ -patch 3 mutant (in C), bmCENP-T^894-1016^ patch 1.1 mutant with the substitutions K895A, R896A, and K907A (in D) and bmCENP-T^894-1016^ patch 1.2 mutant with the substitutions R908S and K911A (in E). Lane L: 100 bp DNA marker. (G) The structure of human CCAN reported in Yatskevitch et al., (2022) highlighting the proximity of α1 of CENP-T HFD (CENPT is coloured brown) to DNA (grey). The side chains of the positively charged amino acids in this region are shown in blue. The illustration was adapted from PDB 7R5S.

**Figure S4.**
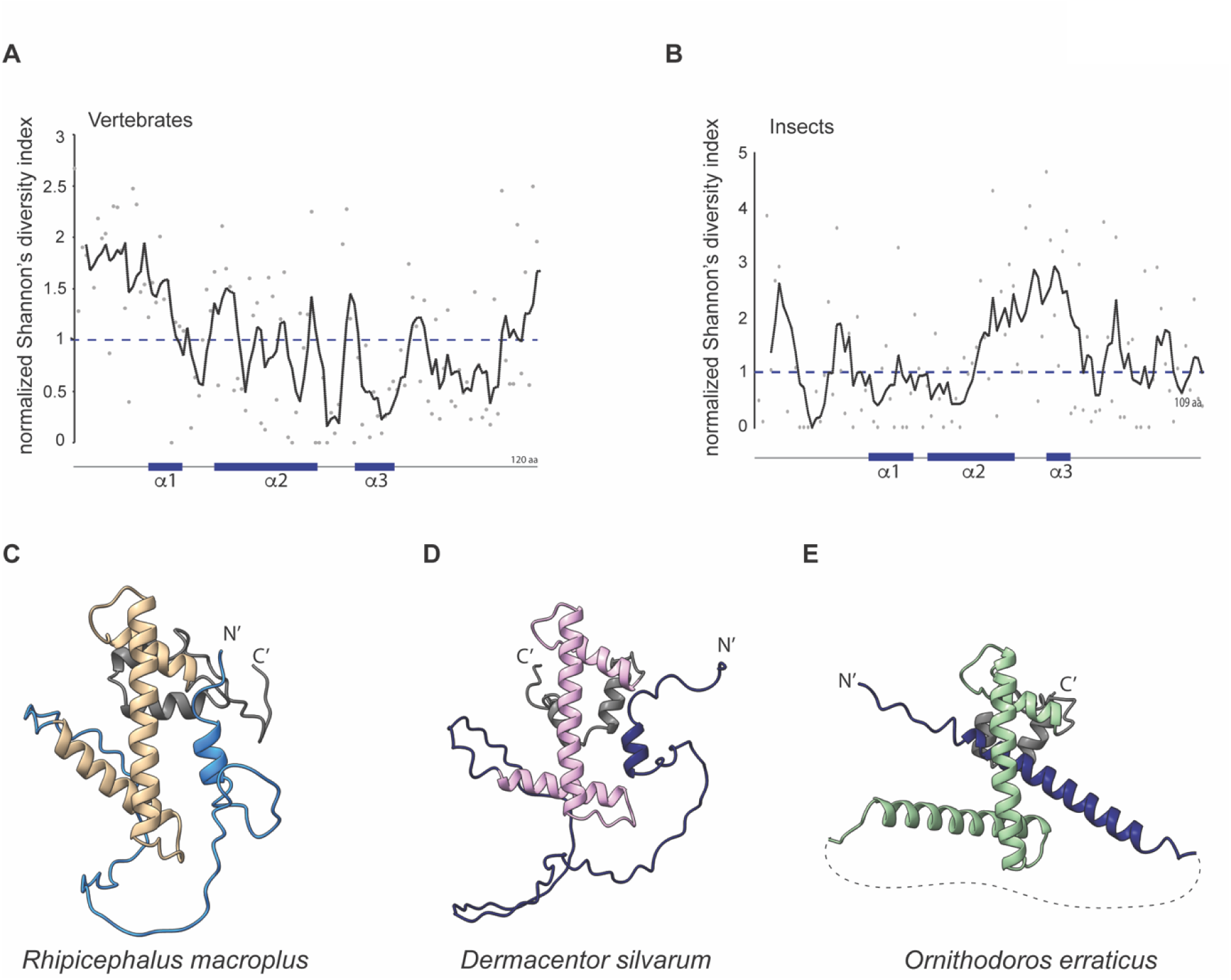
Diversity in the sequence and structure of the cryptic HFD in Arthropoda. (A-B) Comparative analysis of sequence diversity in vertebrate and insect CENP-T using Shannon’s entropy analysis. The grey dots represent the normalized Shannon’s diversity ratio for each amino acid in the C’ of CENP-T. The trendline represents the moving average of 5 datapoints. (C-E) The AlphaFold model for CENP-T from three acariformes in which CENP-W remains undetected. The canonical HFD structure in each of the three species is highlighted in beige, pink, and green respectively. The HFE is coloured grey and a part of the N’ of the protein that blocks the interaction space for CENP-W is shaded blue. For protein IDs see Table S2.

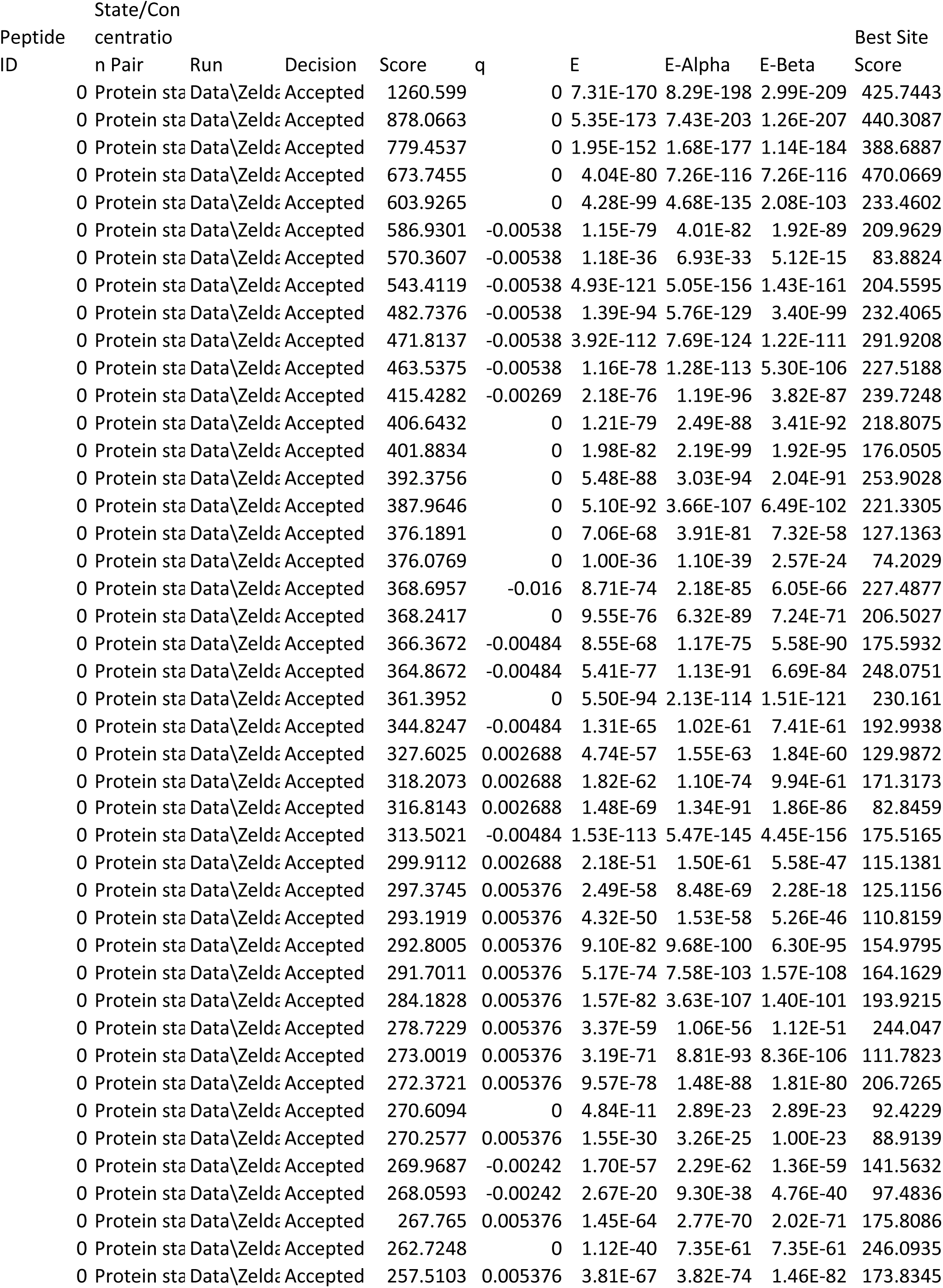
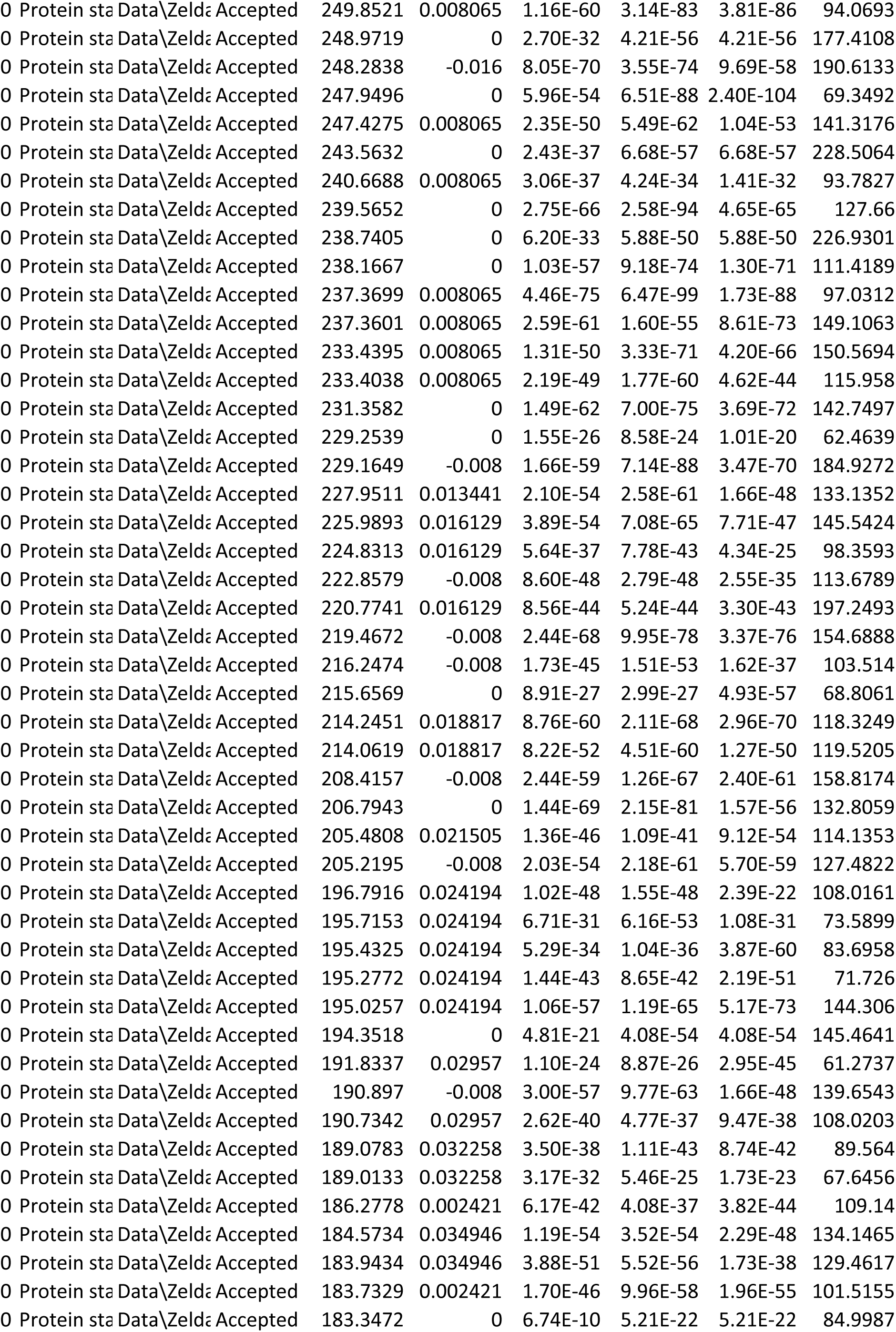
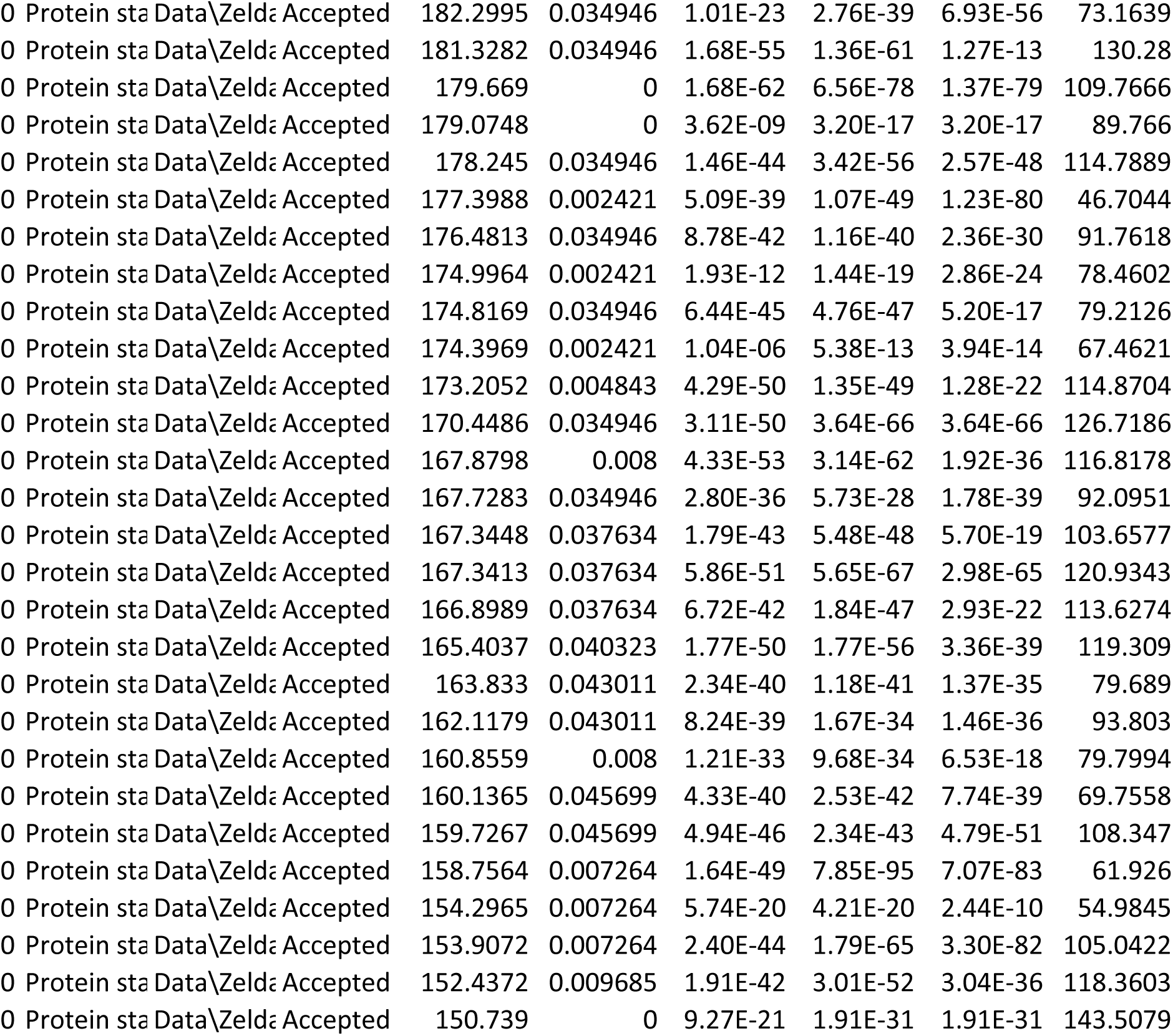

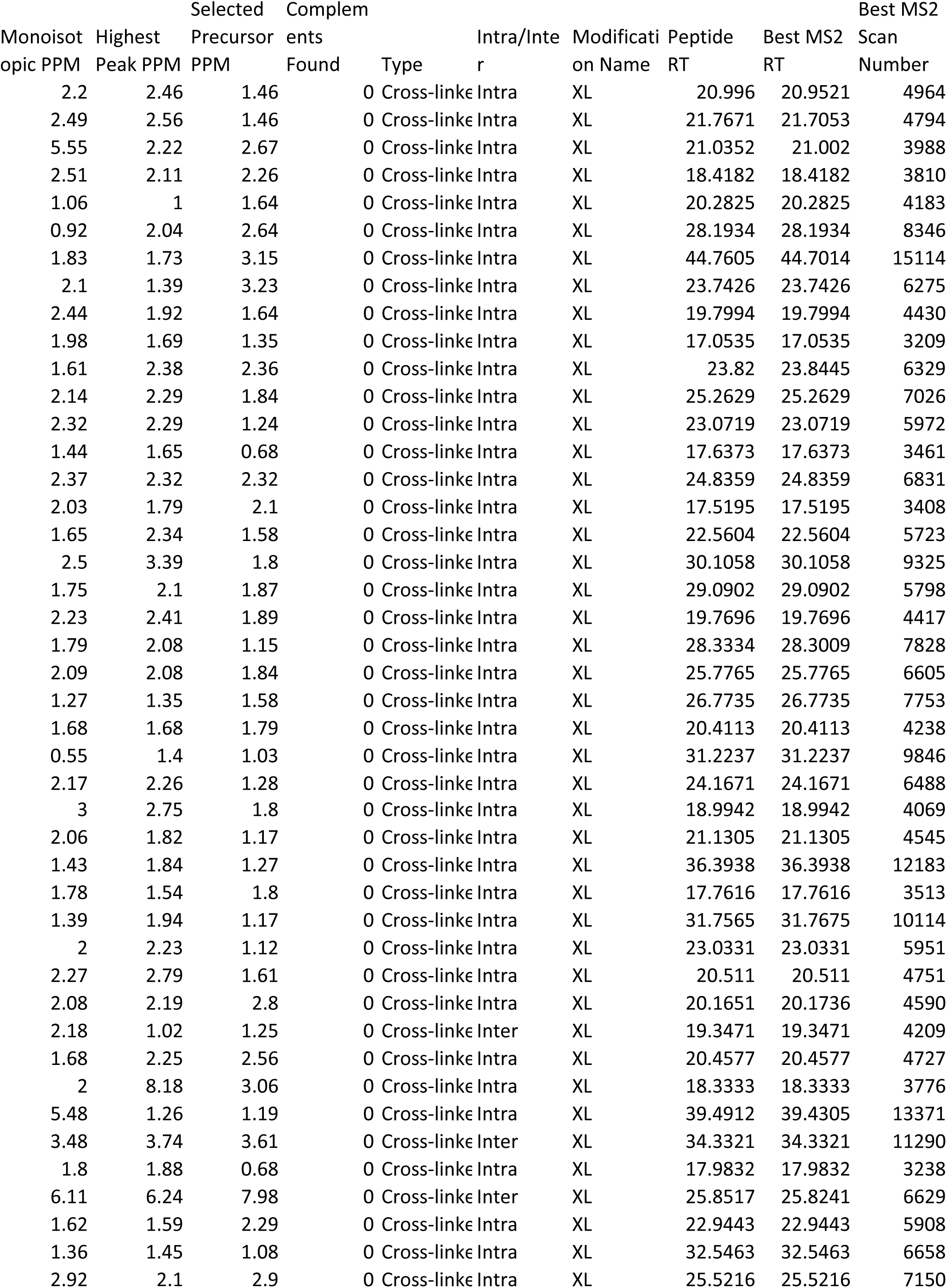
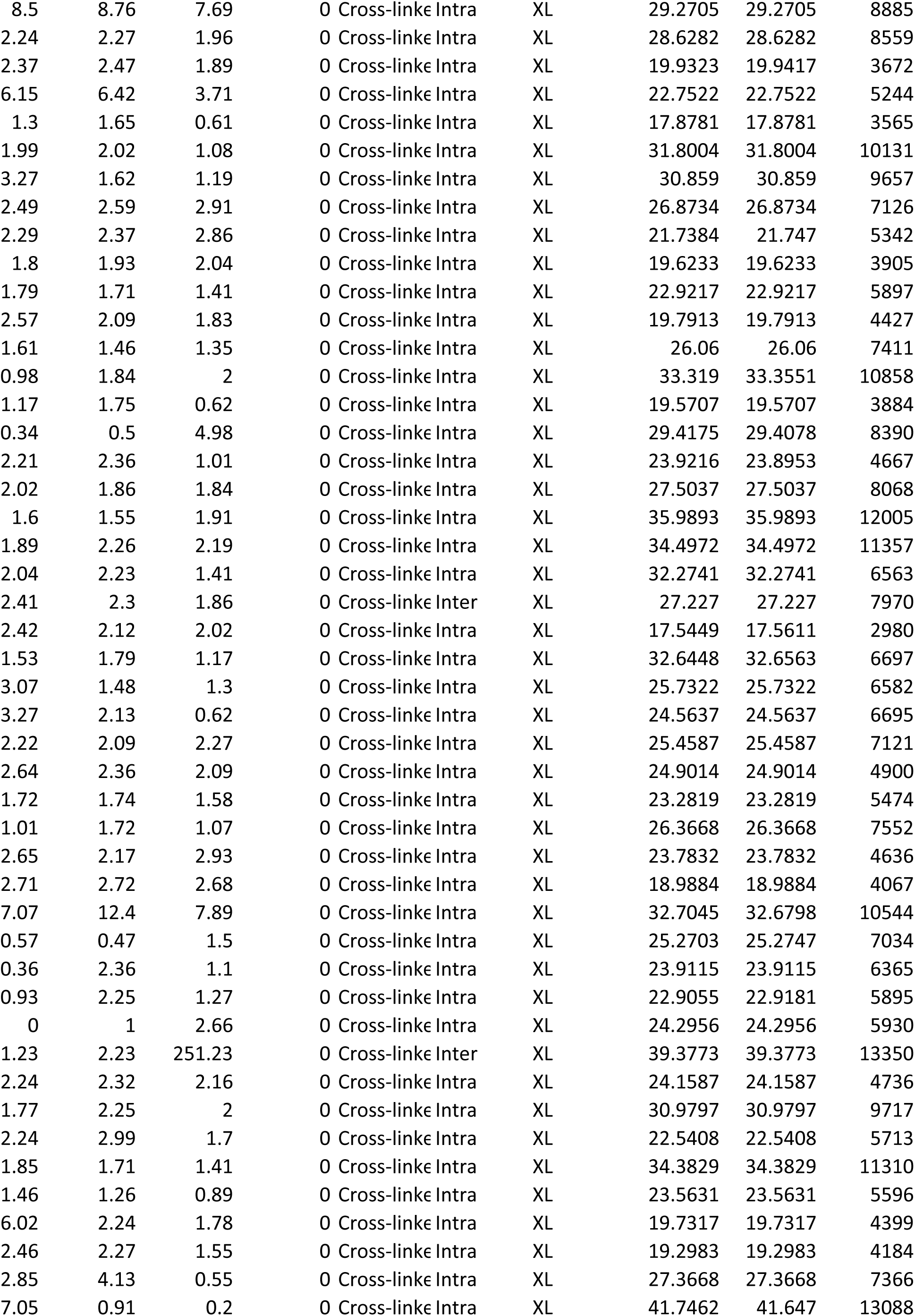
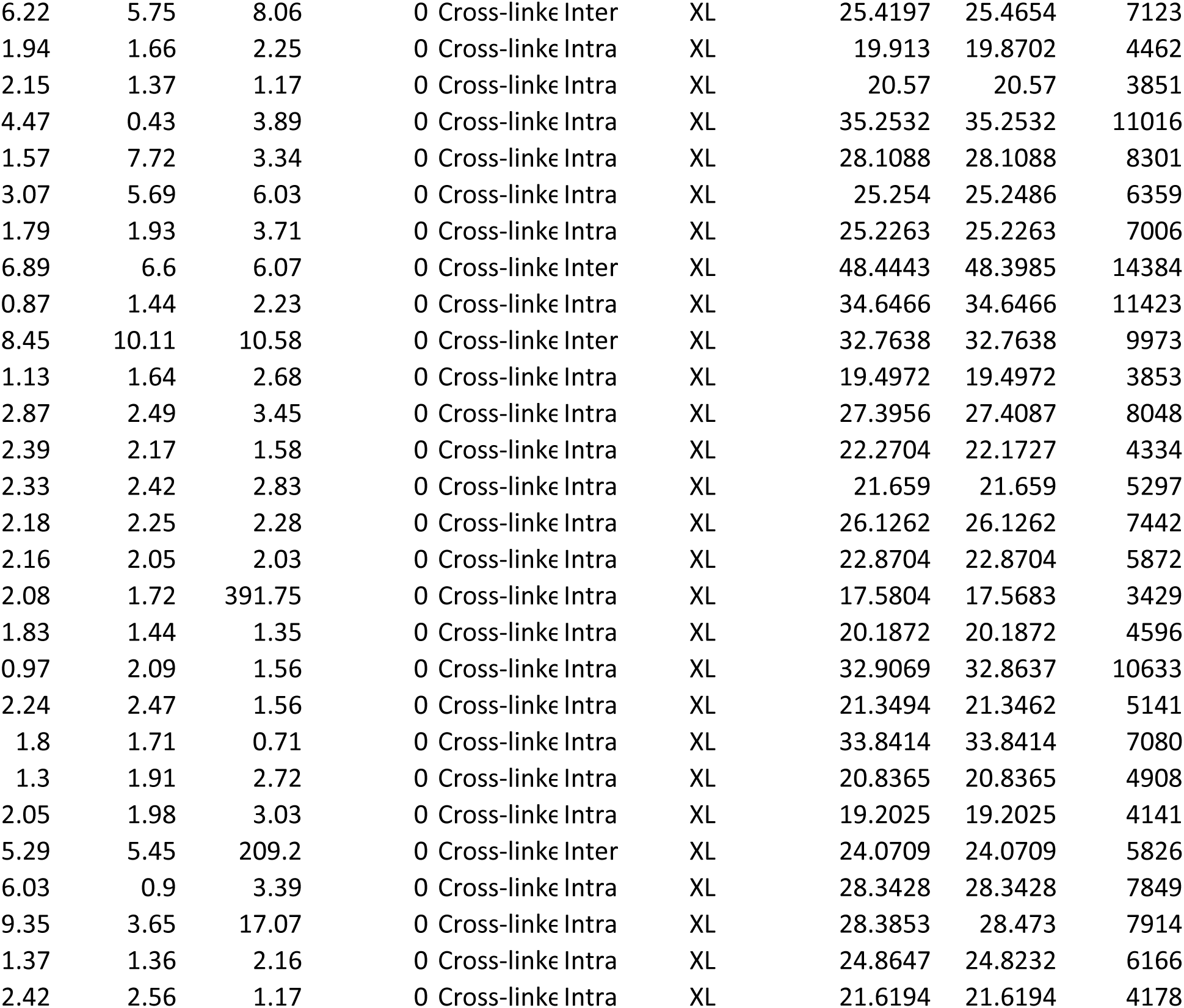

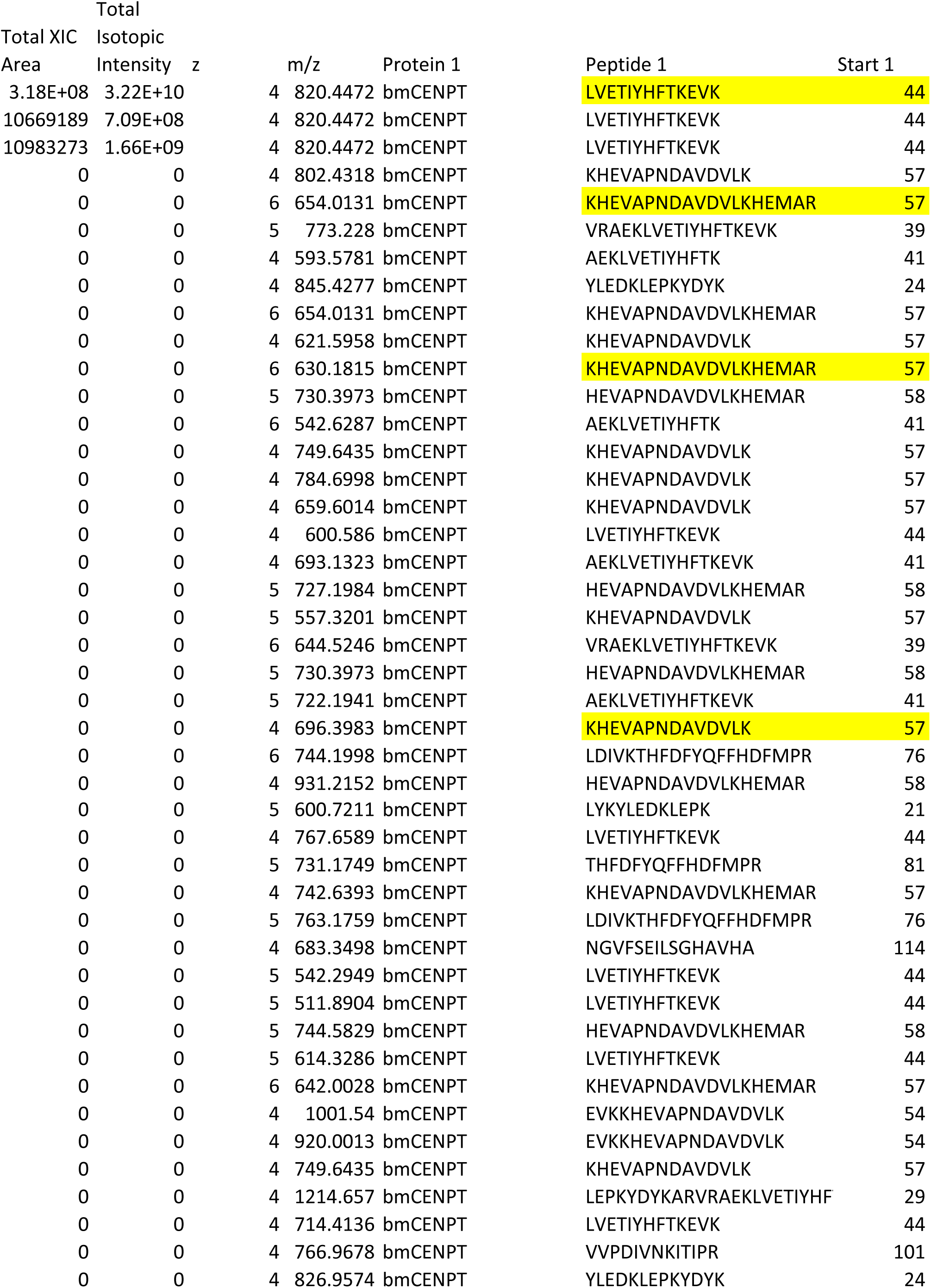
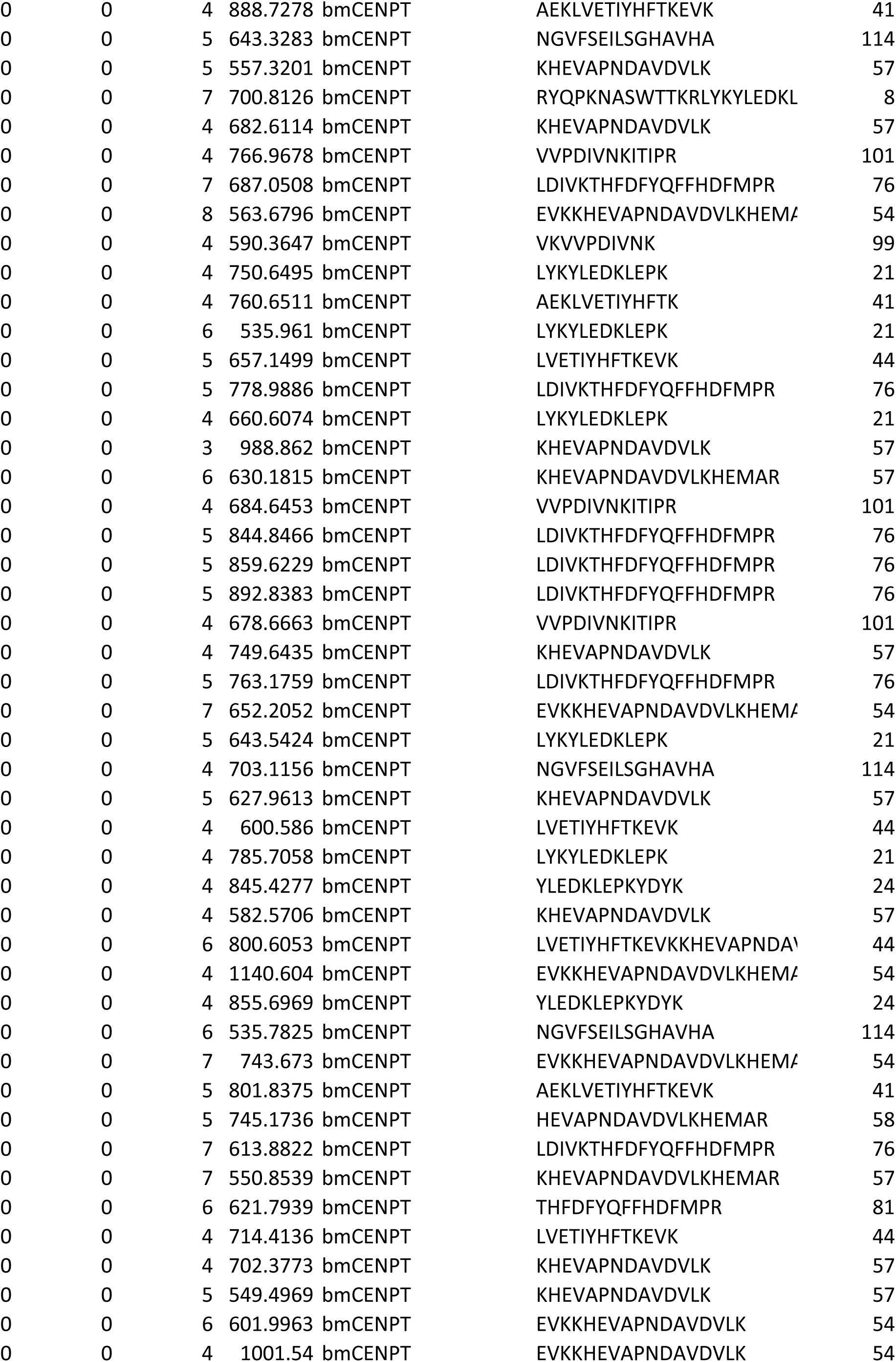
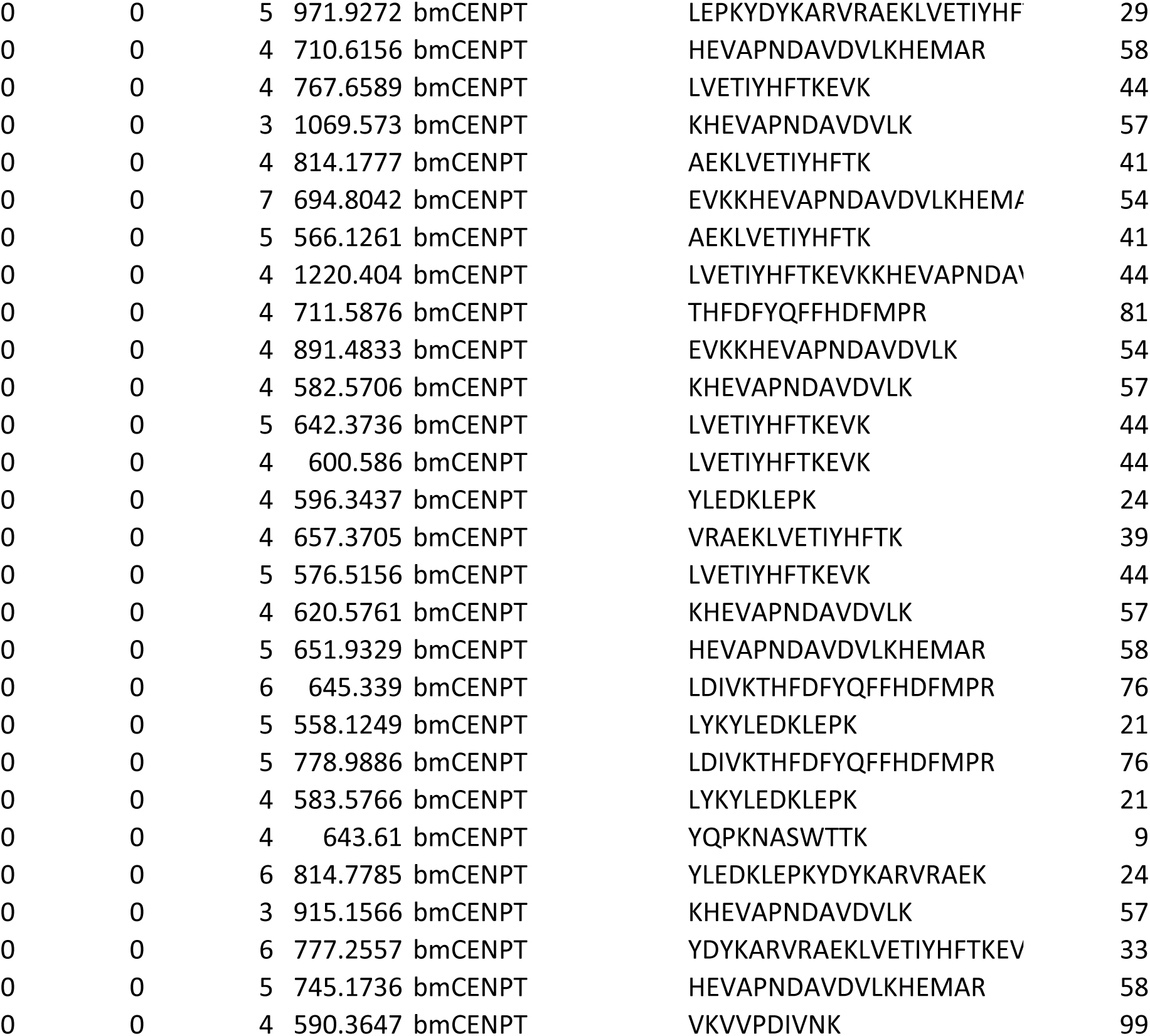

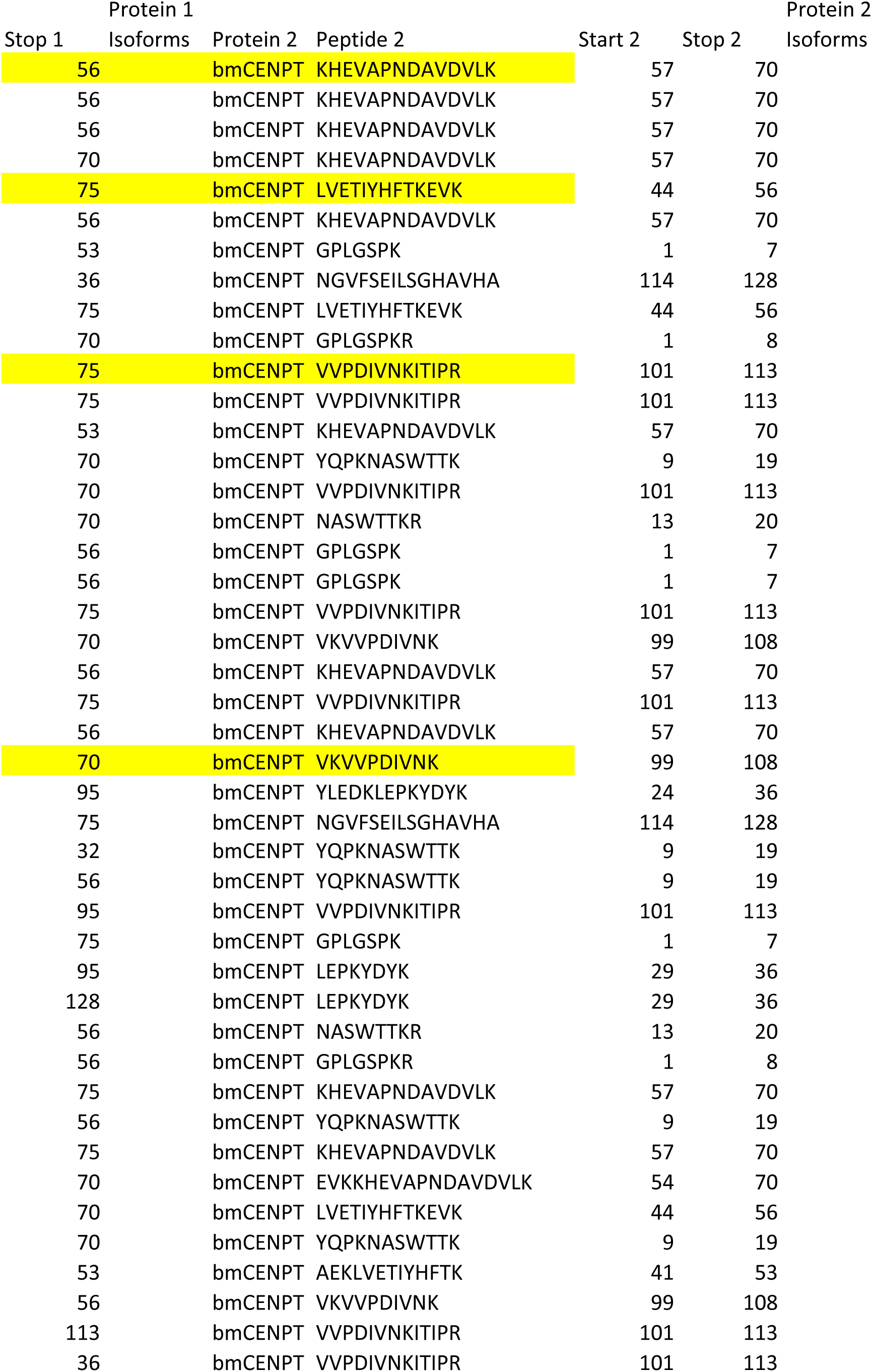
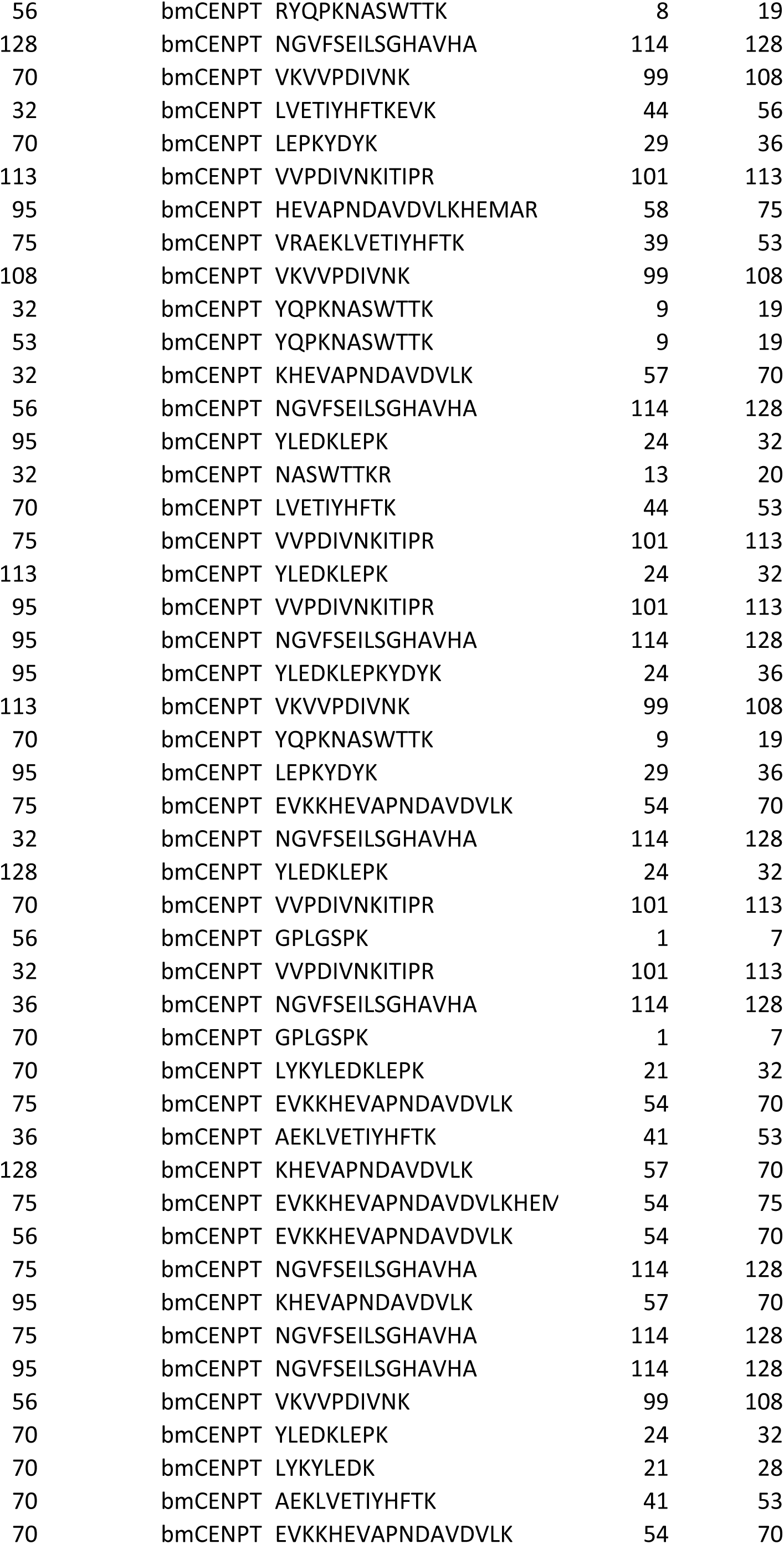
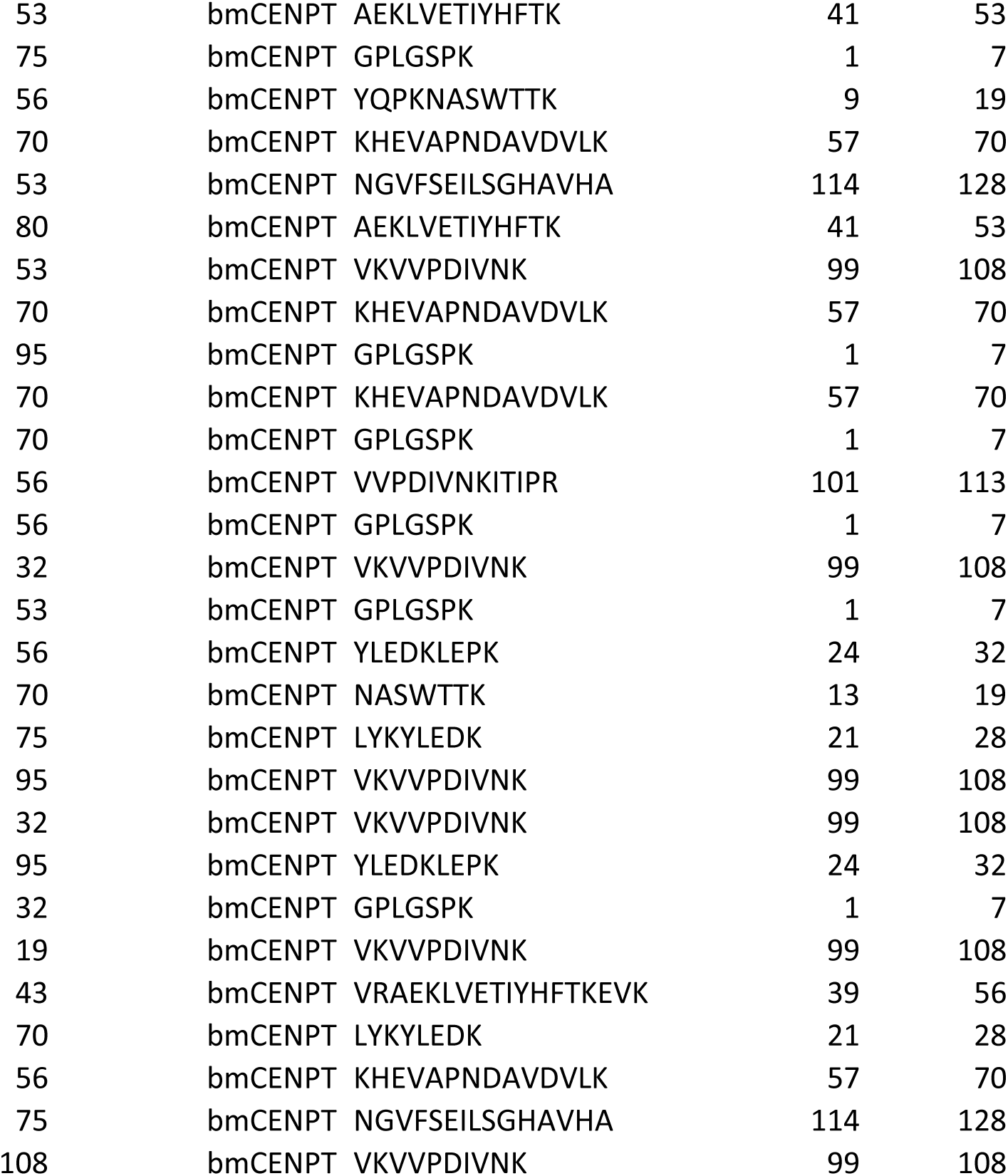

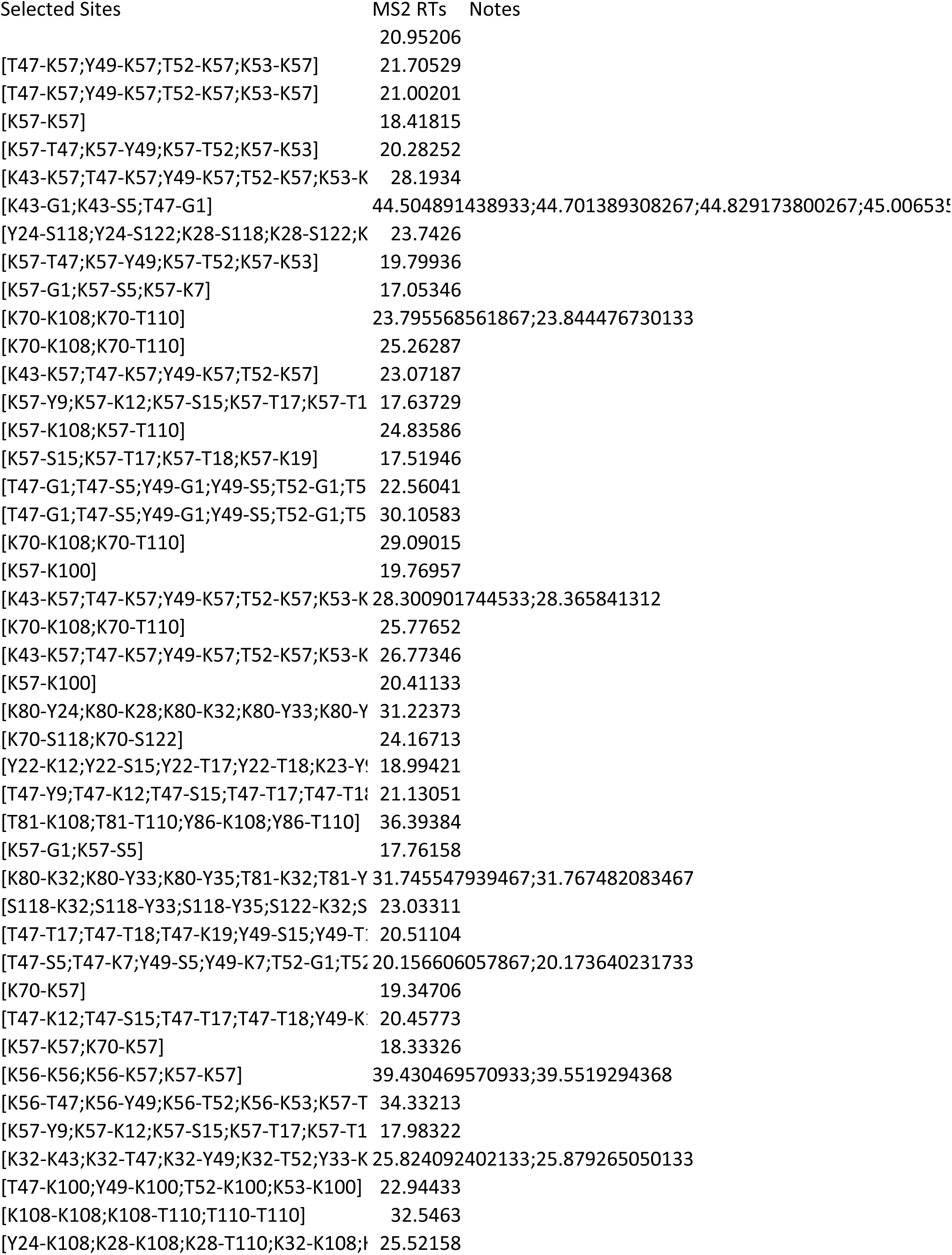
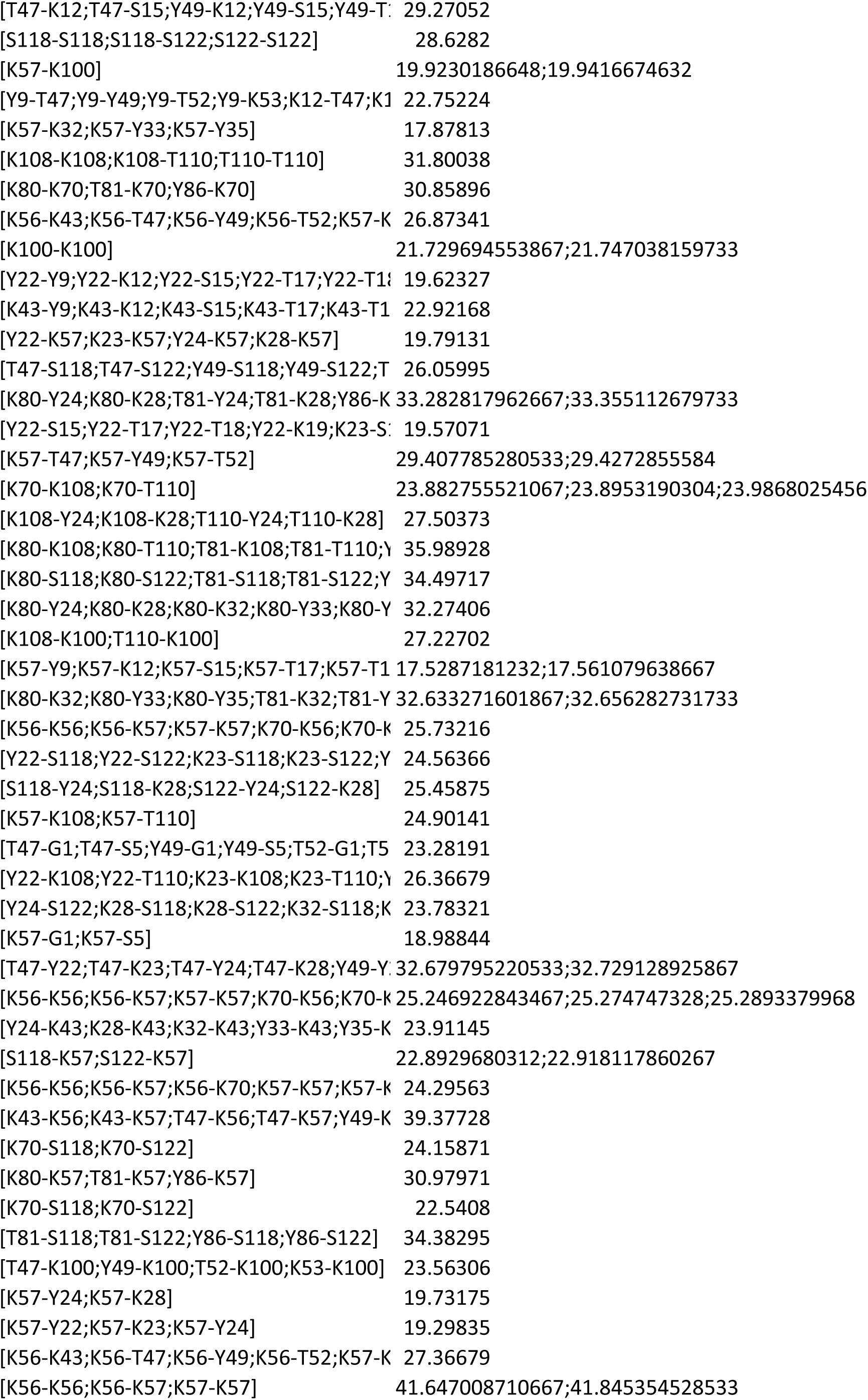
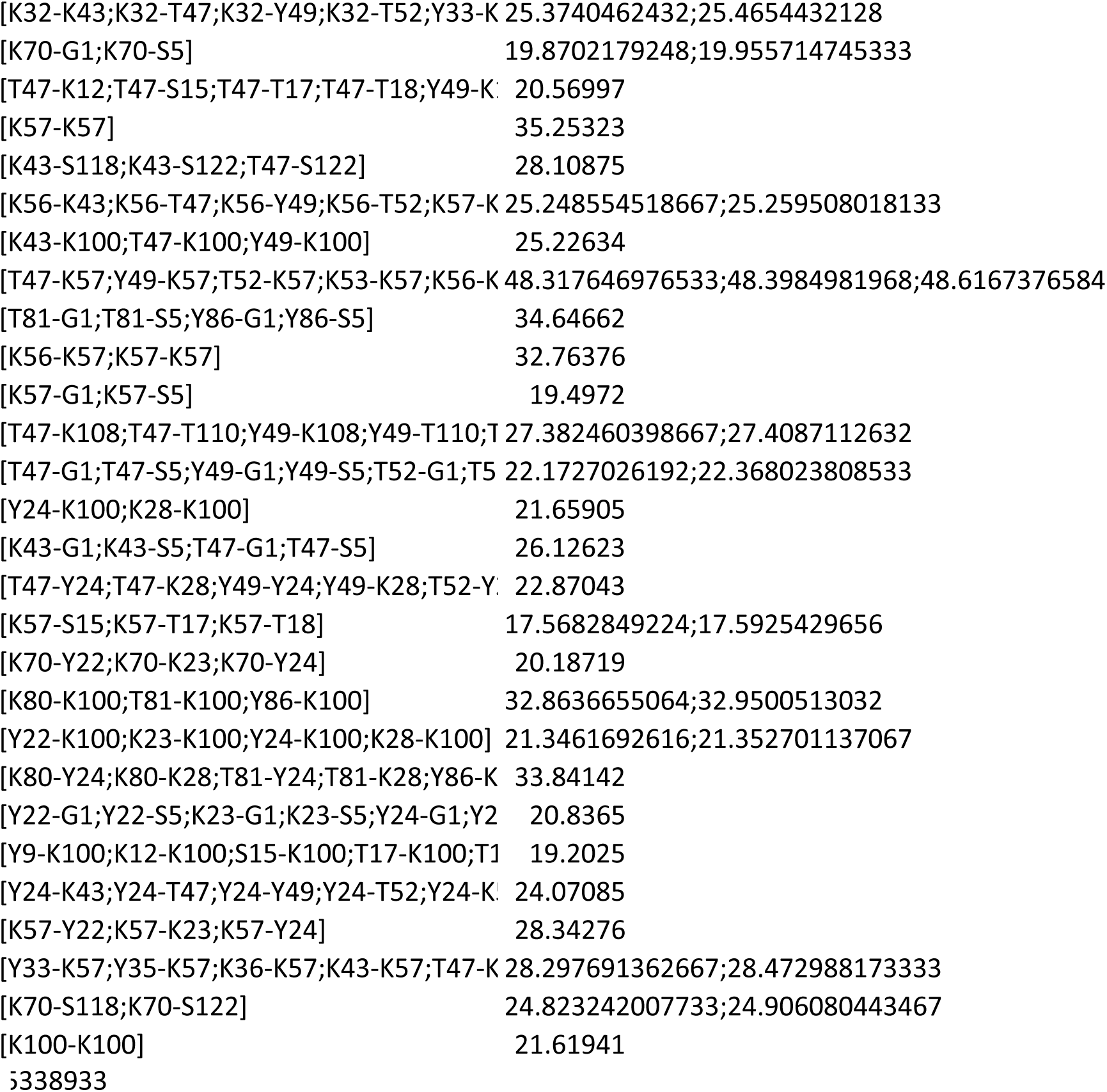

| Organism | CENP-T ID |
| --- | --- |
| *Bombyx mori* | KWMTBOMO06797 |
| *Danaus plexippus* | DPOGS214803-PA |
| *Apis mellifera* | XP_006566152.2 |
| *Bombus impatiens* | XP_024226863.2 |
| *Rhodnius prolixus* | RPRC011181-PA |
| *Gerris buenoi* | KZ651411.1 |
| *Pediculus humanus corporis* | XP_002430294.1 |
| *Blatella germanica* | A0A2P8Y885 |
| *Zootermopsis nevadensis* | A0A067R0W1 |
| *Forficula auricularia* | contig11558 |
| *Bacillus rossius redtenbacheri* | XP_063222590.1 |
| *Acheta domesticus* | NODE_896_length_5051_cov_5.03547_ID_1265 |
| *Libellula vibrans* | NODE_29307_length_498_cov_1.94409_ID_58753 |
| *Ephemera danica* | KZ498339.1:64763-65611 |
| *Peneaus vannameia* | XP_027236760.1 |
| *Helicorthomorpha holstii* | Hho_012284-T1 |
| *Rhipicephalus zambesiensis* | A0A224Y4D1 |
| *Rhipicephalus microplus* | XP_037286617.1 |
| *Dermacentor silvarum* | XP_049524110.1 |
| *Ornithodoros erraticus* | A0A293LVR6 |
| *Limulus polyphemus* | XP_022239208.1 |

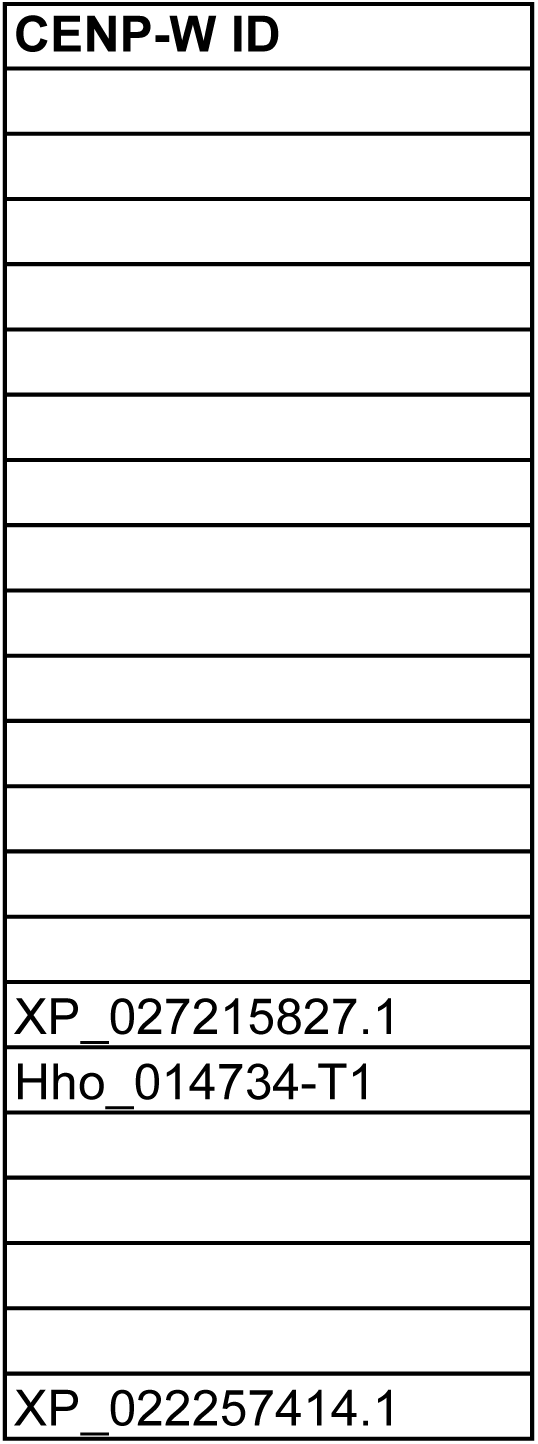

|  |
| --- |
| **Database** |
| http://silkbase.ab.a.u-tokyo.ac.jp/cgi-bin/download.cgi |
| http://reppertlab.org/monarchbase/ |
| http://hymenopteragenome.org/beebase/ |
| http://hymenopteragenome.org/beebase/ |
| https://www.vectorbase.org/organisms/rhodnius-prolixus |
| Cortes-Silva et al, Current Biology 2020 |
| JCVI_LOUSE_1.0 |
| Uniprot |
| Uniprot |
| Cortes-Silva et al, Current Biology 2020 |
| Brsri_v3 |
| Cortes-Silva et al, Current Biology 2020 |
| Cortes-Silva et al, Current Biology 2020 |
| Edan_2.0 |
| NCBI- https://www.ncbi.nlm.nih.gov/bioproject/PRJNA508983 |
| Lok So et al, Nature Communications 2022 (doi: 10.1038/s41467-022-30690-0) |
| Uniprot, https://www.uniprot.org/uniprotkb/A0A224Y4D1/entry |
| NCBI, https://www.ncbi.nlm.nih.gov/bioproject/PRJNA674876 |
| NCBI, https://www.ncbi.nlm.nih.gov/bioproject/PRJNA674874 |
| Uniprot, https://www.uniprot.org/uniprotkb/A0A293LVR6/entry |
| NCBI, https://www.ncbi.nlm.nih.gov/bioproject/PRJNA238073 |

